# Tetracenomycin X sequesters peptidyl-tRNA during translation of QK motifs

**DOI:** 10.1101/2022.08.09.503329

**Authors:** Elodie C. Leroy, Thomas N. Perry, Thibaud T. Renault, C. Axel Innis

**Affiliations:** Univ. Bordeaux, Centre National de la Recherche Scientifique, Institut National de la Santé et de la Recherche Médicale, ARNA, UMR 5320, U1212, Institut Européen de Chimie et Biologie, F-33600 Pessac, France

## Abstract

With antibiotic-resistant bacteria threatening our ability to treat common infections, new lead compounds with distinct target binding sites and limited cross-resistance are urgently needed. Natural products that inhibit the bacterial ribosome – a target for more than half of the antibiotics in use today – are a promising source of such leads and have the potential to be developed into potent drugs through structure-guided design. However, because the mechanisms of action of many of these compounds are not well understood, they are often poor candidates for a structure-based approach. Here, we use inverse toeprinting coupled to next-generation sequencing to show that the aromatic polyketide tetracenomycin X (TcmX) primarily inhibits the formation of a peptide bond between an incoming aminoacyl-tRNA and a terminal Gln-Lys (QK) motif in the nascent polypeptide. Using cryogenic electron microscopy, we reveal that translation inhibition at QK motifs occurs via an unusual mechanism involving sequestration of the 3’ adenosine of peptidyl-tRNA^Lys^ in the drug-occupied nascent polypeptide exit tunnel of the ribosome. Our study provides mechanistic insights into the mode of action of TcmX on the bacterial ribosome and suggests a path forward for the development of novel antibiotics based on a common aromatic polyketide scaffold.

## INTRODUCTION

Aromatic polyketides are a large group of secondary metabolites that include various antimicrobial or antitumor agents, such as tetracycline or doxorubicin, respectively, or the lesser known tetracenomycins^1–4^. Of these, TcmX (Fig. 1a) has been shown to have moderate antimicrobial activity against drug-resistant pathogenic bacteria, including methicillin-resistant *Staphylococcus aureus* and vancomycin-resistant enterococci^5^, and some degree of cytotoxic activity against human cell lines^5–7^. Unlike doxorubicin, with whom it shares a planar tetracyclic core, TcmX does not trigger the SOS response and does not inhibit DNA synthesis in living cells^6^. Like tetracycline^8^, however, TcmX inhibits protein synthesis in bacteria and in human lysates^6^, suggesting that the ribosome and protein synthesis are the physiological targets for this drug. Accordingly, mutating residues U2586, U2609 and U1782 of the *Escherichia coli* (*E. coli*) 23S ribosomal RNA (rRNA) confers resistance to TcmX^6^. Moreover, high-resolution cryogenic electron microscopy (cryo-EM) structures of TcmX bound to the prokaryotic 70S and eukaryotic 80S ribosomes show that the tetracyclic chromophore of TcmX stacks onto the non-canonical base pair formed by 23S rRNA residues U2586 and U1782 (*E. coli* numbering) within the nascent polypeptide exit tunnel^6^. The proximity of this binding site to that of macrolide antibiotics and the observation that, like these sequence-dependent translation inhibitors^9,10^, TcmX only partially obstructs the exit tunnel, suggest that this drug is also likely to block protein synthesis in a manner that depends on the amino acid sequence of the nascent protein^6^. Some evidence in support of this hypothesis comes from the observation that ribosomes translating *ermBL* or *trpL* leader sequences in the presence of TcmX undergo sequence-dependent stalling on Leu-Lys (LK) motifs^6^. However, a comprehensive analysis is needed to assess the generality of these findings, determine the precise mechanism by which TcmX inhibits translation, and provide a molecular basis for the development of TcmX derivatives that potently and specifically inhibit bacterial protein synthesis.

**Fig. 1.**
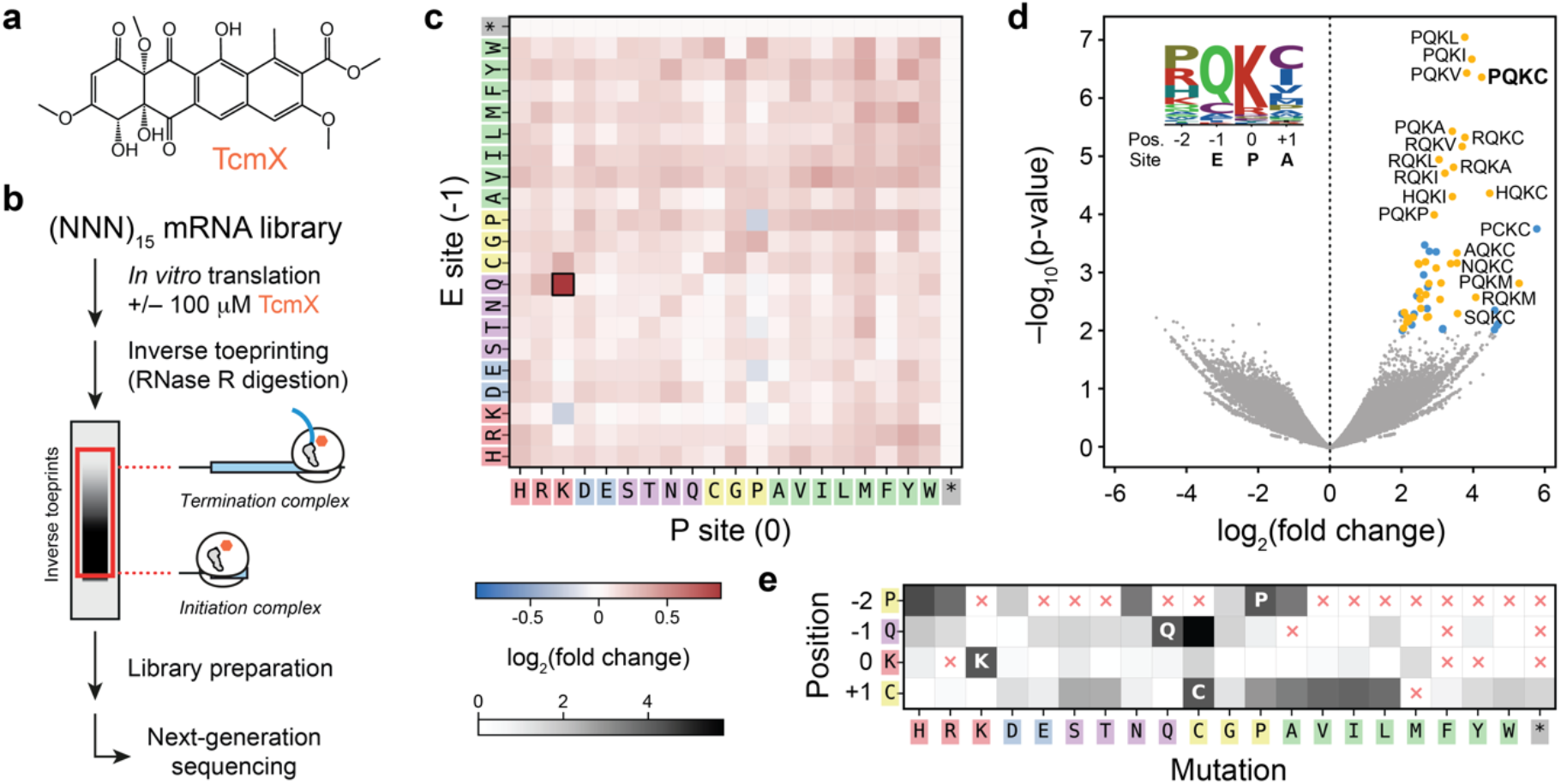
iTP-seq analysis of the context dependence of ribosome inhibition by TcmX. **a**, Chemical structure of TcmX. **b**, Overview of iTP-seq, with a schematic showing the range of inverse toeprints retained after gel electrophoresis. **c**, Heatmap showing the enrichment of motifs at position [-1,0] (E and P sites) after treatment with TcmX. Enrichment is defined as log_2_(*F*_*TcmX*_/*F*_*untreated*_), where *F*_*TcmX*_ is the mean frequency of occurrence of a given motif in the samples treated with TcmX and *F*_*untreated*_ is its mean frequency in the untreated sample. **d**, Volcano plot of statistical significance (-log_10_(p-value)) against enrichment (log_2_(fold change)) for four-amino acid motifs at position [-2,+1]. The majority of highly enriched and statistically significant motifs (top right corner) feature a QK motif at position [-1,0] (yellow points), though a few other motifs were also found (blue points). Thresholds of log_2_(fold change) and -log_10_(p-value) of 2 and 2, respectively, were used to highlight such motifs. **e**, Analysis of single amino acid variants of the PQKC motif, where log_2_(fold change) in motif frequencies relative to the original motif are shown using a gray scale. Red crosses denote enriched variants with a low probability (log_2_(fold change) > 2), but -log_10_(p-value) < 3). The values shown in c, d and e are the mean from three independent experiments.

## RESULTS

### TcmX inhibits translation at QK motifs

To determine how TcmX alters the translational pausing landscape of *E. coli* ribosomes *in vitro*, we used inverse toeprinting coupled to next-generation sequencing (iTP-seq)^11^, a profiling method that locates ribosomes on a library of mRNAs with codon resolution (Fig. 1b). In brief, the position of the leading ribosome on each transcript can be determined by its ability to protect the latter from complete degradation by RNase R, a highly processive RNA exonuclease that digests mRNAs from their 3’ end. Ribosome-protected fragments are amplified to yield a DNA library for next generation sequencing, and the position of leading ribosomes is precisely deduced from the site of RNase R cleavage. Since the entire coding sequence up to the point of ribosome stalling is protected from RNase R digestion, iTP-seq requires no prior knowledge of the sequences expressed. As a result, we could simultaneously monitor the translation of ∼10^12^ distinct short open reading frames, each featuring a stretch of 15 random codons (Extended Data Fig. 1). Translation was carried out using a reconstituted PURE translation system^12^ in the absence or presence of 100 μM TcmX. Following iTP-seq, we measured the frequency of amino acid motifs of various lengths in samples treated with antibiotic and in untreated samples and calculated the fold change in frequency of each motif after antibiotic treatment. For consistency, we shall define the position of all residues/codons in this work relative to the ribosomal P site (position 0), and motifs will be defined according to the range of positions they cover, for example [-1,+1] for a three-amino acid motif spanning the E, P and A sites of the ribosome. When individual positions were considered, we did not observe a significant change in the frequency of amino acids between positions -2 and +1, indicating that TcmX has no detectable impact on the translation of individual codons (Extended Data Fig. 2). In contrast, we could observe a TcmX-dependent 1.7-fold change in frequency for a Gln-Lys (QK) motif at the [-1,0] position, corresponding to stalled ribosomes with a Gln codon in the E site and a Lys codon in the P site (Fig. 1c). To a lesser extent, we also observed a 1.3-fold enrichment in Pro-Gln (PQ) at the [-2,-1] position, and a 1.6-fold enrichment in Lys-Cys (KC) at the [0,+1] position following drug treatment (Extended Data Fig. 2). To our surprise, sequences containing an LK motif at the [0,+1] position did not cause ribosomes to undergo substantial TcmX-dependent stalling in our iTP-seq experiments, as reported in an earlier study^6^ (Extended Data Fig. 2, Extended Data Fig. 3). This may indicate that LK must form part of a longer arrest motif, with neighboring residues within the nascent polypeptide helping to promote ribosome stalling, as observed previously for polyproline motifs^13^. Thus, our iTP-seq results show that TcmX is a context-dependent inhibitor of bacterial translation that causes ribosomes to stall primarily at QK motifs.

To assess whether the sequence surrounding the QK motif affects its ability to induce TcmX-dependent ribosome stalling, we calculated the fold change in the frequencies of all measured four-amino acid motifs following addition of the drug. As expected, nearly all the motifs that caused ribosomes to undergo strong translational arrest contained a QK motif at the [-1,0] position (Fig. 1d). Since PQKC was the most enriched four-amino acid motif with a low p-value (>20-fold change; p=4.4×10^−7^) (Fig. 1d), we estimated the contribution of each of its constituent amino acids to stalling by analyzing all point variants of the motif present in our iTP-seq data set. While most substitutions strongly decreased stalling efficiency, positively charged residues (H,R,K) were tolerated at the -2 position, cysteine at the -1 position, and medium-sized hydrophobic (A,V,I,L) or, to a lesser extent, polar (S,T) residues at the +1 position (Fig. 1d, Fig.1e, Extended Data Fig. 4a). Analysis of XQKX and PXXC variants further showed that residues immediately surrounding the QK motif can be mutated to various extents without compromising ribosome stalling, while mutation of the QK motif itself abolishes stalling – PCKC being a notable exception (Extended Data Fig. 4b-d). Importantly, synonymous codons were enriched to similar extents, suggesting that the nascent amino acid sequence, rather than the mRNA or tRNA sequence, was responsible for the context-dependent response to TcmX (Extended Data Fig. 5). Using classical toeprinting^14,15^, we then confirmed that 100 μM TcmX induces ribosome stalling during translation of an mRNA encoding a MAAAPQKCAAA peptide when tRNA^Lys^ is in the P site of the ribosome, but does not prevent translation of a consecutive stretch of alanine codons (Fig. 2a). Together, these observations reveal the importance of the nascent amino acid sequence surrounding the core QK motif for TcmX-dependent ribosome stalling.

**Fig. 2.**
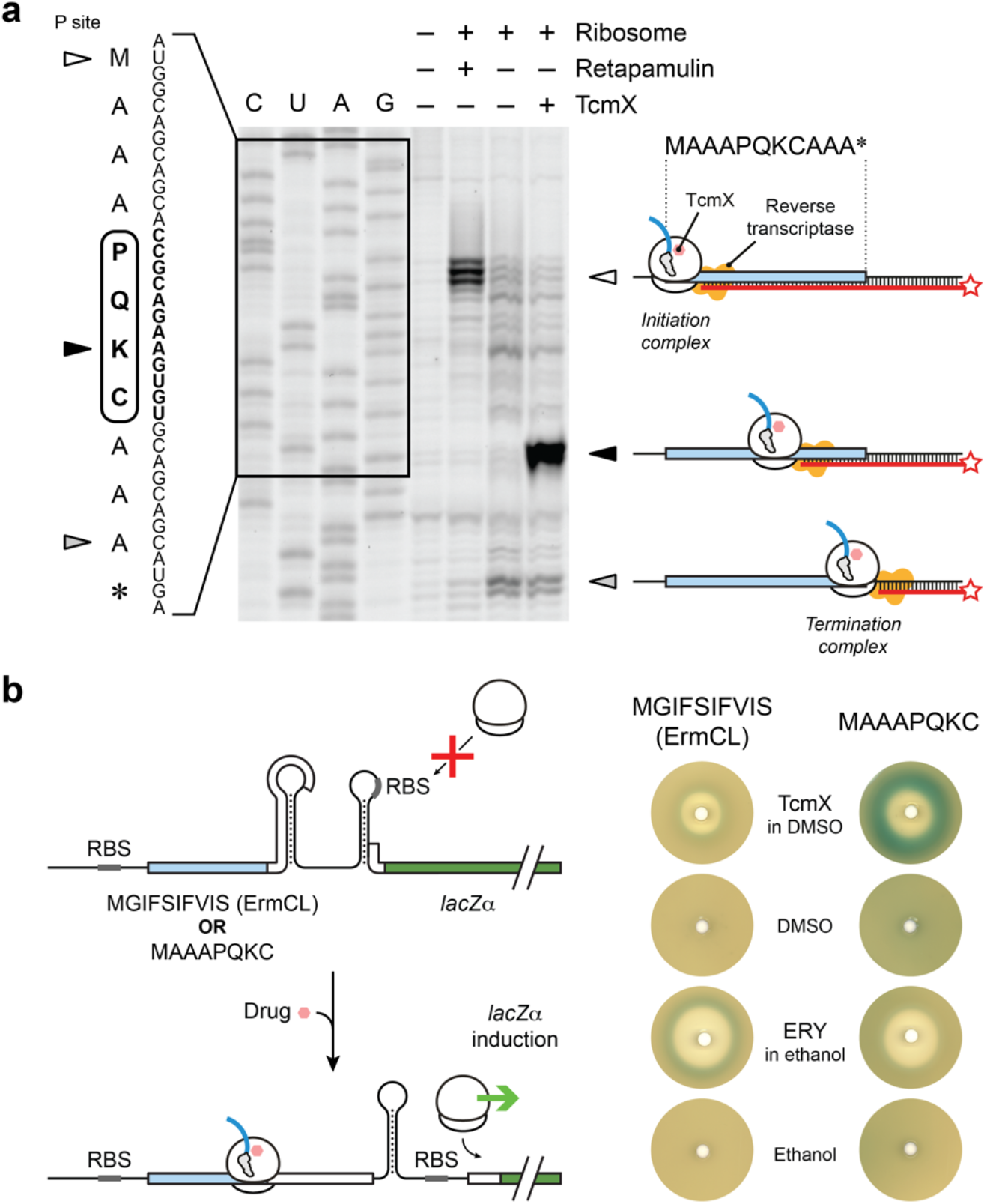
*In vitro* and *in vivo* validation of context-dependent ribosome stalling induced by TcmX. **a**, Toeprinting assay showing the TcmX-dependent stalling of ribosomes translating a MAAAPQKCAAA* sequence in the presence of 100 μM TcmX. Stalling occurred when the K codon was positioned in the P site (black arrow). The antibiotic retapamulin (200 μM) was used as a control to indicate the position of ribosomes stalled on the start codon (white arrow). Ribosomes that reached the stop codon also gave rise to a toeprint corresponding to a termination complex (gray arrow). This experiment was repeated three times independently with similar results. **b**, β-galactosidase complementation assay to monitor *lacZα* expression as a proxy for ribosome stalling. Blue halos result from *lacZα* expression induced by drug-dependent ribosome stalling on the MGIFSIFVIS (*ErmCL*_*1-10*_) or MAAAPQKC leader ORFs. This experiment was repeated four times independently with similar results.

### TcmX hinders QK translation *in vivo*

To determine whether TcmX-dependent translational arrest at PQKC motifs occurs *in vivo*, we performed a β-galactosidase complementation assay that monitors *lacZα* expression as a proxy for ribosome stalling (Fig. 2b). In the original form of this assay^16^, expression of *lacZα* is controlled by a well-characterized translation attenuation mechanism involving the antibiotic-dependent stalling of a ribosome translating the leader open reading frame (ORF) *ermCL*^17–19^. In the absence of added antibiotic, cells express *ermCL* constitutively, but *lacZα* is translationally repressed due to sequestration of its ribosome binding site within an mRNA hairpin. Addition of subinhibitory concentrations of the macrolide antibiotic erythromycin (ERY) causes the ribosome translating *ermCL* to stall on the ninth codon of this leader ORF. This, in turn, leads to a rearrangement of the mRNA secondary structure within the *ermCL-lacZα* intergenic region, which frees the *lacZα* ribosome binding site, enabling the production of α-fragment. As previously shown^20^, ERY-induced ribosome stalling on *ermCL* was manifested by the appearance of a blue halo surrounding the filter disk soaked with this drug, whereas no such halo was observed for untreated cells (Fig. 2b). When, instead, TcmX was added to the disc, only a very weak halo was observed, indicating that TcmX treatment resulted in marginal ribosome stalling on *ermCL* (Fig. 2b). Next, we replaced the first ten codons of *ermCL* with a nucleotide sequence coding for a MAAAPQKC peptide, thus preserving the *ermCL-ermC* intergenic region necessary for *lacZα* induction, as well as the expected position of the TcmX-dependent stalling site observed by toeprinting. When cells carrying this construct were left untreated or were treated with sub-inhibitory concentrations of ERY, no *lacZα* induction was observed (Fig. 2b). However, treatment with TcmX resulted in the appearance of a strong blue halo in the disc diffusion assay, indicating that this antibiotic impacts MAAAPQKC translation, resulting in *lacZα* expression (Fig. 2b). In light of our iTP-seq and classical toeprinting data, these results strongly suggest that the context-dependent action of TcmX on QK motifs observed *in vitro* also occurs *in vivo*.

### TcmX sequesters the 3’ end of P-tRNA

To understand the structural basis of TcmX-dependent translation inhibition at QK motifs, we determined the cryo-EM structure of an *E. coli* 70S ribosome stalled during translation of the PQKC-containing toeprinting template in the presence of 100 μM TcmX, at an overall resolution of 2.7 Å (Fig. 3a, Extended Data Fig. 6). The structure, which we will refer to as 70S-MAAAPQKC-TcmX, was obtained from a major class of particles (48.1%) containing tRNA^Cys^ and tRNA^Lys^ in the ribosomal A and P sites, respectively, as confirmed by the presence of tRNA-specific modifications (Extended Data Fig. 7). A less clearly defined and seemingly aminoacylated tRNA (modeled as tRNA^Gln2^) was also present in the E site, indicating that deacylated tRNA^Gln2^ must first have dissociated from and subsequently been replaced with aminoacyl-tRNA in the ribosomal E site during sample preparation (Extended Data Fig. 7). Local resolution extending to 2.67 Å in the core of the particle enabled us to accurately model all the major ligands including TcmX, peptidyl-tRNA^Lys^ and aminoacyl-tRNA^Cys^ (Fig. 3b, Extended Data Fig. 7). As previously reported^6^, TcmX was bound to the nascent polypeptide exit tunnel (Fig. 3a,b), where it stacks against the noncanonical U2586:U1782 base pair (Extended Data Fig. 8a,b). In the A site, we could observe clear density for a cysteinyl moiety attached to tRNA^Cys^ (residue C^+1^) (Fig. 3b). The bases of 23S rRNA residues U2506, G2583 and U2584 were in their induced conformation, but 23S rRNA residue U2585 and residue A76 of the P-site tRNA remained in the uninduced conformation^21^, indicating only partial aminoacyl-tRNA accommodation into the peptidyl transferase center (Extended Data Fig. 9a-d). In the P site, clear density was also visible for all but the first two residues of the nascent peptide (A^-4^A^-3^P^-2^Q^-1^K^0^) (Fig. 3b), confirming that the last amino acid incorporated into the nascent peptide was a lysine residue, in agreement with our iTP-seq and toeprinting data.

**Fig. 3.**
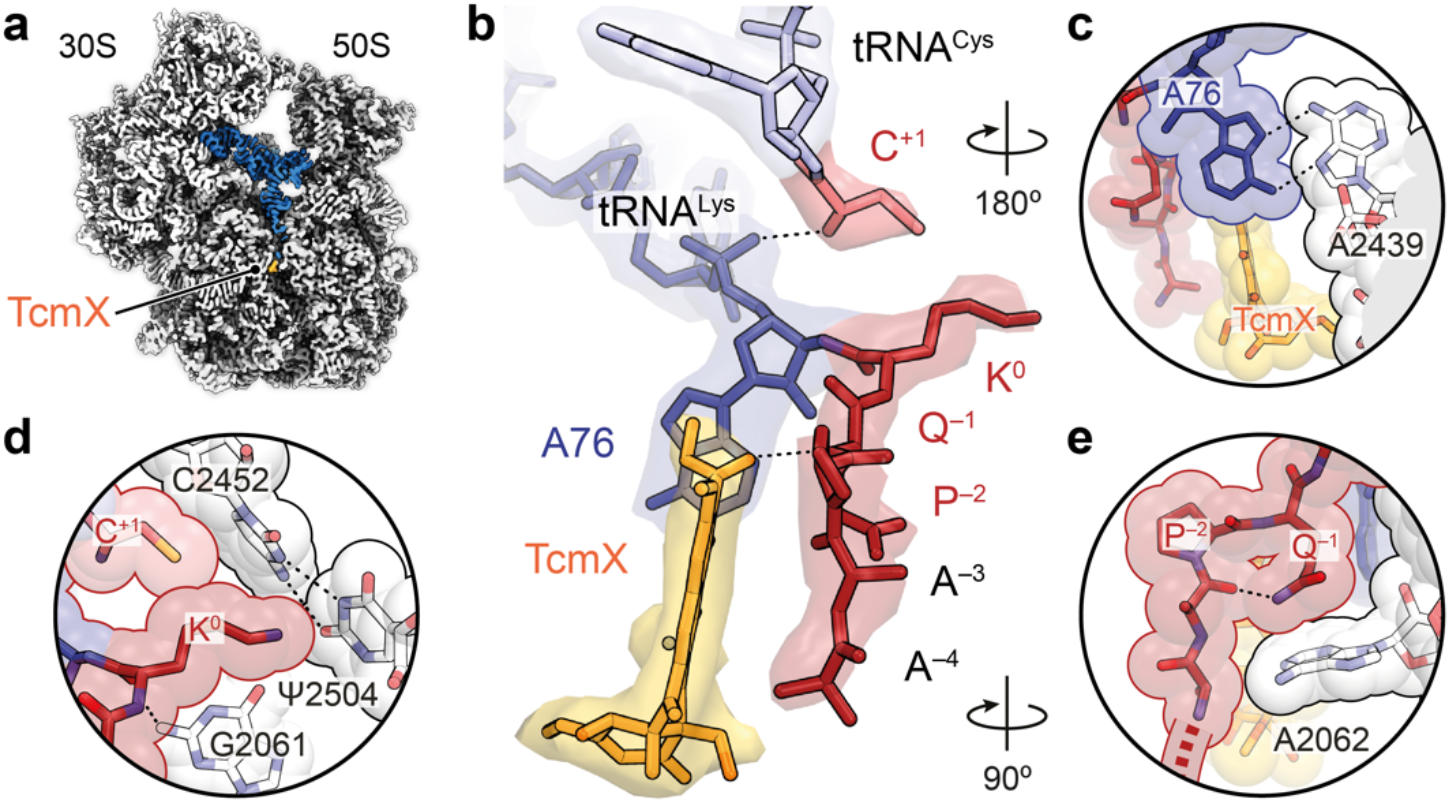
Cryo-EM structure of a 70S-MAAAPQKC-TcmX complex. **a**, Transverse section of the cryo-EM map for the 70S-MAAAPQKC-TcmX complex with isolated densities showing the ribosome (light gray), peptidyl-tRNA (blue) and TcmX (orange). **b**, Molecular model and surrounding map densities for the MAAAPQK nascent chain (red), P-site tRNA (dark blue), A-site tRNA (light blue), incoming residue C^+1^ (salmon) and TcmX (orange). **c**, Residue A76 of the P-site tRNA forms a non-canonical *trans* Hoogsteen–Hoogsteen base pair with residue A2439 of the 23S rRNA. **d**, Residue K^0^ from the nascent peptide occupies the A-site crevice, where it is likely to interfere with the accommodation of some but not all aminoacyl-tRNAs. **e**, The side chain of residue Q^-1^ from the nascent peptide stacks against the base of 23S rRNA residue A2062. It also forms a hydrogen bond with the backbone carbonyl oxygen of residue -3, which is likely to be facilitated by the kink induced in the nascent peptide by residue P^-2^.

Unexpectedly, residue A76 of the P-site tRNA was rotated and shifted by nearly 12 Å compared to the pre-attack state, in which peptidyl-tRNA and aminoacyl-tRNA are poised to undergo peptide bond formation^22^. This atypical conformation of residue A76 is distinct from those observed previously in structures of the ribosome in complex with antibiotics that perturb the position of the 3’ end of peptidyl-tRNA, such as blasticidin S^23,24^, or aminoacyl-tRNA, such as hygromycin A or A201A^25^ (Extended Data Fig. 8c-f), and causes the base of 23S rRNA residue A2602 to stack against the repositioned N-terminus of ribosomal protein bL27 (Extended Data Fig. 9e). Several elements stabilize the highly unusual peptidyl-tRNA conformation induced by TcmX, most notably a *trans* Hoogsteen-Hoogsteen base pair between residue A76 and 23S rRNA residue A2439 (Fig. 3c). On the peptide side, residue K^0^ occupies the A-site crevice, where its backbone carbonyl oxygen forms a hydrogen bond with the amine of 23S rRNA residue G2061 (Fig. 3d). The side chain of residue Q^-1^ stacks against the base of 23S rRNA residue A2062 (Fig. 3e). Mutating the residue at position -1 to the shorter asparagine would likely result in suboptimal stacking with A2062, explaining why this conservative substitution is not enriched in our iTP-seq dataset. The side chain and backbone amide groups of residue Q^-1^ also form intramolecular hydrogen bonds with the backbone carbonyl of residue A^-3^ (Fig. 3e) and with TcmX (Fig. 3b), respectively. Together with residue P^-2^, whose presence is not strictly required at this position but which may nevertheless contribute to stalling efficiency (Fig. 1d,e), these interactions help to stabilize a kink within the nascent peptide that promotes favorable van der Waals contacts with the drug (Fig. 3b) and the ribosome (Fig. 3e). The notable absence of amino acid sequence-dependent hydrogen bonds or salt bridges between the QK-containing peptide and the drug-bound ribosome suggests that the sequence specificity of the interaction could lie in this Q^-1^- dependent kink and in the interaction of the K^0^ side chain with the A-site crevice, both of which contribute to ensuring that peptidyl-tRNA featuring a QK motif at the [-1,0] position packs optimally against the ribosome and TcmX. In summary, the structure of the 70S-MAAAPQKC-TcmX complex observed by cryo-EM features a highly disrupted peptidyl transferase center stabilized by a network of interactions between the drug, the QK-containing peptidyl-tRNA and the ribosome.

### Mechanism of ribosome inhibition by TcmX

Under physiological conditions, the α-amino group of an incoming aminoacyl-tRNA is deprotonated to yield an –NH_2_ nucleophile capable of attacking the ester bond connecting the C-terminal amino acid of the nascent peptide to the P-site tRNA. For peptide bond formation to occur, the attacking nucleophile must be located within a relatively short distance (∼3 Å) of the carbonyl ester carbon (Fig. 4a)^21^. In our structure, the distance between the α-amine of the incoming cysteine (C^+1^) and the carbonyl ester carbon of the terminal lysine (K^0^) of the nascent peptide is increased to 5.2 Å, and the geometry of the reactants is highly unfavorable for peptidyl transfer (Fig. 4b). In addition, the α-amine of residue C^+1^ appears to form a salt bridge with the backbone phosphate of tRNA^Lys^ residue A76 in the P site, suggesting that its protonation state (– NH_3_^+^) renders it unable to perform a nucleophilic attack (Fig. 4b). The poor placement of the reactants and the lack of a –NH_2_ nucleophile resulting from the likely protonation of the α-amino group are thus likely to account for the inability of the ribosome to form a peptide bond after QK motifs in the presence of TcmX.

**Fig. 4.**
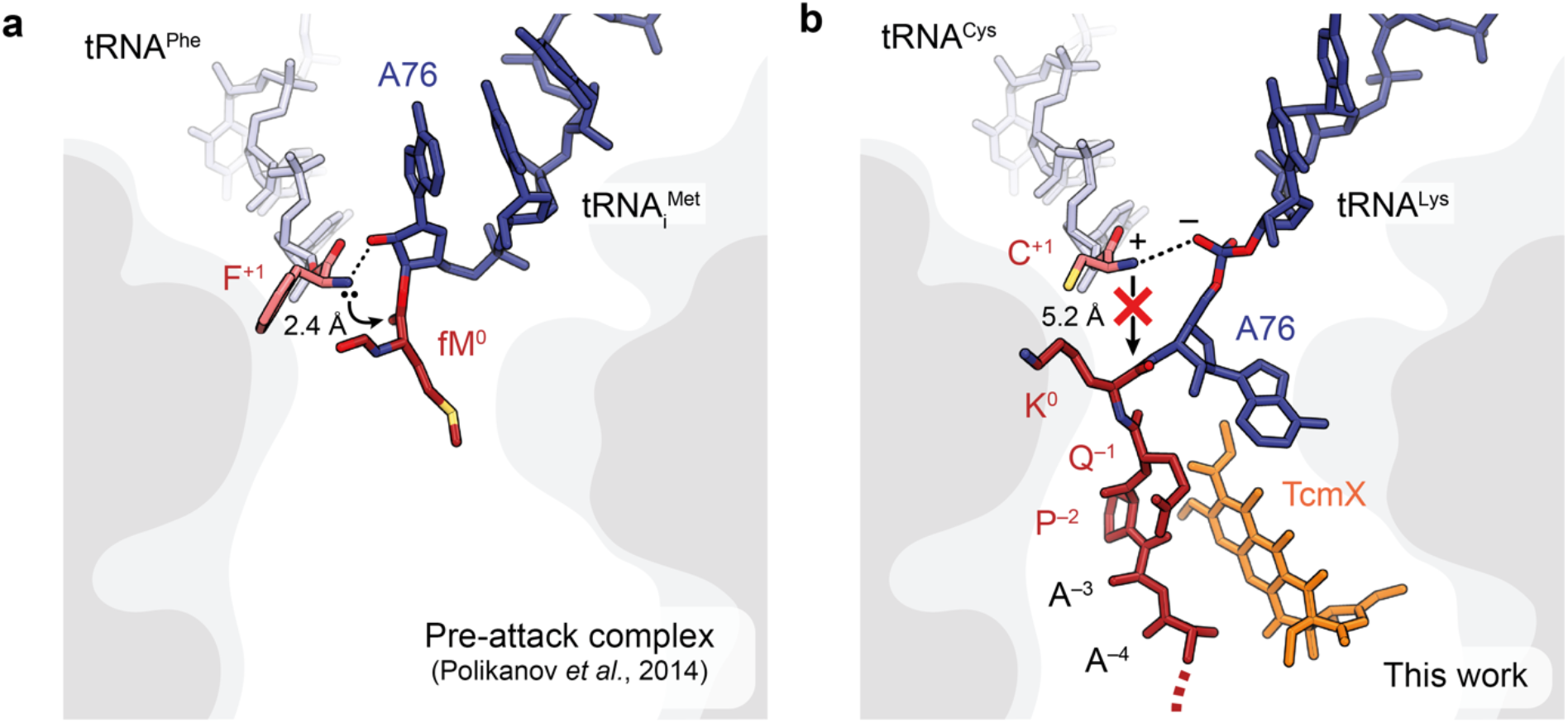
Mechanism of TcmX-dependent translation inhibition at QK motifs. **a**, Pre-attack complex (PDB 1VY4)^22^ where phenylalanyl-tRNA^Phe^ in the A site is poised to attack formyl-methionyl-tRNA ^Met^ in the P site. This complex, which was obtained using a non-reactive form of formyl-methionyl-tRNA ^Met^ containing a non-hydrolyzable amide bond instead of the usual ester bond, features an attacking nucleophile positioned 2.4 Å away from the ester carbonyl carbon. **b**, In the 70S-MAAAPQKC-TcmX complex, sequestration of residue A76 of the P-site tRNA into the nascent polypeptide exit tunnel of the ribosome increases the distance between the attacking amine and the ester bond to 5.2 Å, making peptide bond formation less likely. Moreover, a salt bridge between the attacking amine and the backbone phosphate of P-site tRNA residue A76 renders protonation of the α-amino group highly likely.

## CONCLUSION

In this work, we reveal the principal mechanism by which the recently discovered antibiotic TcmX inhibits bacterial translation upon binding to the nascent polypeptide exit tunnel of the ribosome. By using iTP-seq to characterize the stalling landscape of TcmX-bound *E. coli* ribosomes, we showed that TcmX inhibits translation at QK motifs, in a manner that is dependent on the nascent amino acid sequence. Using complementary biochemical and structural approaches, we identified how a nascent peptide featuring a C-terminal QK motif interacts with the bacterial ribosome and TcmX to inhibit peptide bond formation *in vitro* and *in vivo*.

The mechanism of action of TcmX is reminiscent of that by which macrolide antibiotics inhibit the ribosome^26^. Indeed, both macrolides and TcmX inhibit translation in a peptide sequence-dependent manner. In the presence of macrolides, translational arrest occurs primarily at +X+ motifs, where + represents a positively charged amino acid like arginine or lysine^27^. Both TcmX and macrolides bind near the peptidyl transferase center, albeit on opposite faces of the nascent polypeptide exit tunnel^6,26^, explaining the lack of cross-resistance to TcmX and macrolides^6^. These different binding sites and sequence specificities ultimately result in substantially distinct mechanisms of action. On a macrolide-bound ribosome, nascent peptides containing a +X+ motif adopt a conformation where the first positively charged residue within the motif (typically an arginine at position -1) occupies the A-site crevice and prevents the accommodation of an incoming positively charged amino acid into the A site, leading to ribosome stalling^27^. In contrast, translation of QK motifs in the presence of TcmX appears to only mildly impair aminoacyl-tRNA accommodation, but instead results in the sequestration of the peptidyl-tRNA 3’ end inside the nascent polypeptide exit tunnel, and subsequent inhibition of peptide bond formation.

In order to develop TcmX into a clinically useful drug, the lack of specificity of this antibiotic for bacterial over human translation must first be overcome. In this respect, our findings could guide the structure-guided design of TcmX derivatives that induce ribosome stalling at QK motifs in bacteria while showing reduced binding to the human ribosome. Although the immediate environment of TcmX is similar in the *E. coli* and human ribosomes, a cavity located ∼10 Å away from the O-methyl group of ring D of TcmX in the bacterial ribosome is blocked by the base of 28S rRNA residue U1591 in the human ribosome (Extended Data Fig. 10). Derivatives of TcmX carrying a suitable side chain at this position may be identified that depend on binding to this cavity, resulting in greater affinity and specificity for the bacterial target. Other tetracenomycins, such as elloramycin or tetracenomycin C, exert antimicrobial activity against a variety of *Actinomyces* and *Streptomyces* species^28–32^ and may operate through similar context-dependent mechanisms. Elucidation of the modes of action of this very promising family of natural products could open new avenues for the development of potent antibiotics to treat multidrug-resistant infections.

## METHODS

### General experimental procedures for iTP-seq

iTP-seq was performed as described previously, with modifications^11^. DNA and RNA products at various points of the iTP-seq protocol were analyzed on 9% acrylamide (19:1) TBE (90 mM Tris, 90 mM boric acid, and 2 mM EDTA) gels and stained with SyBR Gold (Invitrogen). Inverse toeprints were excised from 12% acrylamide TBE gels using a clean scalpel. RNA gel electrophoresis was performed under denaturing conditions (8 M urea in the gel). All reactions were performed using molecular biology grade water (Millipore). Reagents and oligonucleotides used in this study are listed in Supplementary Table 1.

### Preparation of an (NNN)_15_ mRNA library for iTP-seq

A DNA library composed of short ORFs with a random N-terminal stretch of 15 NNN (aNy aNy, aNy) codons was purchased from Eurogentec as a single stranded oligonucleotide (*iTP_NNN15_random-library*). An expression cassette for *in vitro* transcription and translation was obtained by polymerase chain reaction (PCR) with Phusion DNA polymerase (6 cycles [98°C, 10 s; 62°C, 5 s; 72°C, 10 s]), using the *iTP_NNN15_random-library* oligonucleotide in combination with oligonucleotides *iTP/TP_frag1_T7_RBS_ATG_f* and *iTP_Stop_EcoRV_r* as templates (0.1 pmol of each oligonucleotide per 50 μl reaction) and oligonucleotides *iTP/TP_frag1_T7_f* and *iTP_EcoRV_r* for amplification (1 pmol of each oligonucleotide per 50 μl reaction). The PCR product was purified using a PCR purification kit (Qiagen) according to the manufacturer’s instructions and quantified with a 2100 Agilent Bioanalyzer. The sequence of the final (NNN)_15_ DNA template was:

CGATCGAATTC<u>TAATACGACTCACTATA</u>GGGCTTAAGTATAAGGAGGAAAAAAT**ATGNNNN NNNNNNNNNNNNNNNNNNNNNNNNNNNNNNNNNNNNNNNNNGCGATCTCGGTGT GA**TGA*GATATC*AATATCAAAAAGGATCCATATA (the T7 promoter is underlined, the translated region is in bold; the EcoRV site for iTP-seq is in italics).

In order to generate an mRNA library for iTP-seq, *in vitro* transcription was performed at 20°C for 3 h, in a 200 μl reaction volume containing 8 pmol of (NNN)_15_ library, 3.7 μl of 20 mg/ml stock of homemade T7 RNA polymerase (Addgene plasmid p6XHis-T7-P266L^33^, 80 mM Tris–HCl pH 7.6, 24 mM MgCl_2_, 2 mM spermidine, 40 mM DTT, 7.5 mM ATP (Sigma Aldrich), CTP and UTP, 0.3 mM GTP (CTP, UTP, and GTP from Jena Bioscience), and 4.5 mM Thio-Phosphate-GMP (Genaxxon Bioscience). Transcripts were purified using an RNA Clean & Concentrator-5 purification kit (Zymo Research) according to the manufacturer’s instructions. The concentration of mRNA was determined using a NanoDrop spectrophotometer (Thermo Fisher Scientific).

As described previously^11^, 5’-biotinylation of the mRNA templates for *in vitro* translation is required to facilitate the various enzymatic reactions performed for iTP-seq. For the biotinylation reaction, biotin-maleimide (Vectorlabs) was dissolved in dimethylformamide according to the manufacturer’s instructions, and 800 pmol mRNA were mixed with 800 nmol biotin-maleimide in 100 mM Bis-Tris-acetate buffer pH 6.7 in a total volume of 50 μl and incubated at room temperature for 2.5 h. Unincorporated biotin was removed by washing the mRNA three times with H_2_O (molecular biology grade, Millipore) in an Amicon membrane centrifugal concentrator with a molecular weight cutoff (MWCO) of 30 kDa (Millipore). mRNA was recovered and the biotinylation efficiency was determined by means of a dot blot, as follows. H+ bond membrane

(GE Healthcare) was treated with 6Å∼ SSC buffer (900 mM NaCl, 90 mM Na_3_-citrate, pH 7.0) for 10 min and dried briefly between two pieces of Whatman paper. Biotinylated (NNN)_15_ mRNA and a 5’-biotinylated oligonucleotide standard (*iTP_Biotin_standard*) were diluted in 6x SSC buffer to 0.5, 1.0, 2.5, and 5.0 μM, and 1 μl of each dilution was pipetted onto the prepared membrane. The membrane was baked for 2 h at 80°C to adsorb nucleic acids, and subsequently blocked in 2.5% dry milk solution in TBS-T (50 mM Tris–HCl, 150 mM NaCl, and 0.05% [vol/vol] Tween-20, pH 7.5) for 1 h at room temperature. The milk solution was removed and the membrane was incubated with a 1:1,000 dilution of streptavidin-alkaline phosphatase antibody (Promega) in TBS-T for 1 h at room temperature. Unbound antibody was removed by washing three times with TBS-T buffer. Colorimetric detection was performed using an NBT/BCIP detection kit (Promega) according to the manufacturer’s instructions. The membrane was imaged immediately on a Bio-Rad Imager. The biotinylation efficiency was estimated by comparing the intensity of the sample dots with the intensity of the standard dots.

Finally, biotinylated mRNA must be polyadenylated at the 3’ end to improve the efficiency of RNase R digestion during inverse toeprinting. Polyadenylation was performed using Poly-A polymerase (New England Biolabs) and the buffer supplied by the manufacturer. The ratio of biotinylated mRNA to ATP molecules was chosen to be 1:100. The reaction was incubated at 37°C for 2 h and the efficiency of the polyadenylation reaction was assessed by denaturing polyacrylamide gel electrophoresis (PAGE) (9%). Polyadenylated mRNA was purified using the RNA Clean & Concentrator-5 kit (Zymo Research) according to the manufacturer’s instructions.

### *In vitro* translation and inverse toeprinting

*In vitro* translation was carried out with a custom PURExpress **Δ**RF-123 **Δ**Ribosome kit (New England Biolabs), using ∼5 pmol of 5’-biotinylated and 3’-polyadenylated mRNA as a template. TcmX dissolved in dimethyl sulfoxide (DMSO) was supplemented at a final concentration of 100 μM in 5 μl reactions. Release factors 1 and 3 were added to the translation reaction according to the manufacturer’s instructions. Translation was performed at 37°C for 30 min, after which the samples were placed on ice and 5 μl ice-cold Mg^2+^ buffer (50 mM Hepes-KOH, 100 mM K-glutamate, 87 mM Mg-acetate, and 1 mM DTT, pH 7.5) was added to the reactions, thereby increasing the Mg^2+^ concentration to 50 mM. 1 μl of RNase R (1 mg/ml) was added, followed by an additional incubation for 30 min at 37°C to ensure complete mRNA degradation. 139 μl of 1x BWT buffer was added to stop the reaction (5 mM Tris–HCl, 0.5 mM EDTA, 1 M NaCl, and 0.05% [vol/vol] Tween-20, pH 7.5).

### Library preparation for next generation sequencing

For each sample, 5 μl of M-280 streptavidin Dynabead (Thermo Fisher Scientific) were washed three times with 1x BWT buffer in DNA loBind tubes (Eppendorf) and resuspended in 50 μl of the same buffer. Dynabeads and RNA from the previous step were combined into these tubes and incubated on a tube rotator for 15 min at room temperature to allow binding of the biotinylated mRNA to the streptavidin beads. After incubation, beads were collected using a magnet and the supernatant was discarded. The beads were washed one time with 1x BWT buffer to remove unbound RNA, followed by two washes with H_2_O to remove the 1x BWT buffer. Beads were resuspended in 9.5 μl of linker ligation reaction mixture containing 4 μl of water, 1 μl of 10x T4 RNA ligase 2 (truncated) buffer (New England Biolabs), 3 μl of 50% PEG 8,000 (New England Biolabs), 1 μl of *iTP_3’_linker_ApoI* (10 μM) and 0.5 μl of ligase (T4 RNA ligase 2, truncated - 200 000 U/ml - New England Biolabs) per reaction. Linker ligation was allowed to proceed on a tube rotator for 2.5 h at room temperature.

Following ligation of the linker, beads were washed once with H_2_O to remove unincorporated linker oligonucleotide and were resuspended in 18.5 μl of reverse transcription reaction mixture containing 11.5 μl of water, 1 μl of dNTPs (10 mM of each – New England Biolabs), 1 μl of *iTP_Linker_r* oligonucleotide (2 μM), 4 μl of first strand synthesis buffer (5X - Thermo Fisher Scientific) and 1 μl of DTT (0.1 M - Thermo Fisher Scientific) per reaction. The samples were incubated for 5 min at 65ºC to anneal the primer to the complementary sequence and then placed on ice. 1 μl of reverse transcriptase (Superscript III – 200,000 U/ml – Thermo Fisher Scientific) was added to each tube and the samples were incubated for 30 min at 55ºC in a Thermomixer at 500 rpm to allow reverse transcription of the Dynabead-bound mRNA.

Reverse transcribed cDNA was used without further purification as a template for PCR. To generate double stranded DNA for restriction digestion, a fill-up reaction was performed using *iTP_cDNA_f* oligonucleotide and the reverse transcribed cDNA (10 s denaturation, 10 s annealing at 42°C, and 30 s elongation at 72°C). The resulting double stranded DNA was combined with 1 μl of EcoRV-HF restriction enzyme and the sample was incubated at 37°C for 1 h. To amplify undigested DNA, *iTP_Linker_r* oligonucleotide was added, and 10-16 cycles of PCR were performed (denaturation at 98°C for 10 s, annealing at 60°C for 10 s, and elongation at 72°C for 10 s). The number of PCR cycles was adjusted to give a visible band on the gel while minimizing non-specific byproducts.

Bands containing inverse toeprints corresponding to stalled ribosomes from the initiation codon to the last NNN codon were excised from the gel with a clean scalpel. Gel pieces were crushed through a 5 ml syringe into 15 ml Falcon tubes and 10 ml of gel elution buffer (10 mM Tris–HCl, pH 8.0, 500 mM Na-Acetate, and 0.5 mM Na-EDTA) were added. The tubes were incubated on a tube rotator at room temperature overnight. Gel debris were separated from the extraction solution by filtering through 0.22 μm centrifugal filters (Millipore). Each sample was then concentrated to ∼1 ml using a SpeedVac. DNA was precipitated in 5 ml Eppendorf tubes using 1 ml of isopropanol with 3.7 μl GlycoBlue (Thermo Fisher Scientific) and incubating at −80°C overnight. After precipitation, DNA was recovered by centrifugation in a ThermoScientific Heraeus Multifuge X3R centrifuge at 20,000xg for 30 min at 4°C using a Fiberlite F15-8×50cy rotor (ThermoScientific). The supernatant was removed, and DNA pellets were resuspended in 20 μl H_2_O (molecular biology grade, Millipore) for subsequent addition of the next-generation sequencing (NGS) adaptors.

Long NGS adaptor oligonucleotides (*iTP_NGS_adaptor_f* and the reverse oligonucleotides *iTP_NGS_adaptor_index[…]*) contain Illumina TruSeq adapter sequences followed by 18 nucleotides complementary to the 5’ or 3’ region of the cDNA. The reverse oligonucleotides also contain barcode sequences for multiplexing according to the TruSeq v1/v2/LT protocol (Illumina). Sequencing libraries were obtained from 12–16 cycles of PCR using 0.02 μM long NGS adapter oligonucleotides (forward and reverse) and 0.2 μM short amplification oligonucleotides (*iTP*_*NGS_f* and *iTP_NGS_r*). PCR products were purified using a Qiagen PCR purification kit. The size and concentration of the fragments obtained were measured using a 2100 Agilent Bioanalyzer with the DNA 1000 kit.

### Next generation sequencing and analysis of iTP-seq data

Next generation sequencing was performed by the BGI Facility in Hong Kong, on an Illumina Hiseq X Ten system in rapid run mode with 150 PE reads.

Unless indicated otherwise, data analysis was carried out using a series of custom Python scripts. Read pairs were assembled using PEAR v0.9.10^34^, with the maximal proportion of uncalled bases in a read set to 0 (**–**u option) and the upper bound for the resulting quality score set to 126 (**–**c option). The 5′ flanking region was defined as GTATAAGGAGGAAAAAAT and the 3′ flanking region was GGTATCTCGGTGTGACTG. A maximum of two mismatches within each of these flanking regions was tolerated, whereas all other reads were discarded. Trimming of the retained reads resulted in sequences with a start codon directly at the 5′ end and the site of RNase R cleavage at the 3′ end (Extended Data Fig. 1). Reads were filtered using a custom script (parse_filter_fastq.py) to have a minimum quality score of 30 at all positions, contain no Ns, and be devoid of aberrations (*e*.*g*. known contaminants or abnormally long stretches of A) and inverse toeprints were aligned relative to the ribosomal P site. The data underlying the heatmaps and volcano plots were then analyzed using another custom script (make_report.py) to count the number of reads per motif for various combinations of positions, combined with DESeq2^35^ to compute the enrichment statistics. The graphs were produced with Matplotlib 3.3.2^36^. The logo of the top arrest motifs was generated with Logomaker^37^.

### Toeprinting assays

Toeprinting was performed as described previously^15^. Briefly, a DNA template containing a T7 promoter, a ribosome binding site, a sequence coding for the MAAAPQKCAAA peptide and the NV1 sequence^19^ was generated by PCR using as templates oligonucleotides *iTP/TP_frag1_T7_RBS_ATG_f, TP_frag2_r, TP_frag2_NV1_r* and *TP_MAAAPQKCAAA*_f* (0.1 pmol of each oligonucleotide per 50 μl reaction), and the short primers *iTP/TP_frag1_T7_f* and *TP_short_r* for amplification (1 pmol of each oligonucleotide per 50 μl reaction). The sequence of the *TP_ MAAAPQKCAAA\** toeprinting mRNA template was:

CGATCGAATTC<u>TAATACGACTCACTATA</u>GGGCTTAAGTATAAGGAGGAAAAAAT**ATGGCAG CAGCACCGCAGAAGTGTGCAGCAGCATGA***AGCGAATAATAACTGACTCTGAACAACATCC GTACTCTTCGTGCGCAGGCAAGG*TTAATAAGCAAAATTCATTATAACC (the T7 promoter is underlined; the coding region is in bold; the NV1 sequence is in italics).

DNA templates were transcribed and translated *in vitro* using a custom PURExpress ΔRF-123 ΔRibosome Kit (New England Biolabs) and a final concentration of 2.2 μM ribosomes. Ligands were dissolved in water and added as needed at the beginning of the reaction. The Yakima Yellow-labelled probe complementary to the NV1 sequence (*TP_RT_yakima-yellow_r*) was added to the 5 μl reaction after incubating for 15 min at 37°C (2 μM) and the sample was incubated for another 5 min at the same temperature. Reverse transcription was then performed with 50 U of Avian Myeloblastosis Virus reverse transcriptase (Promega Corporation) for 20 min at 37 °C. RNA was degraded by adding 0.5 μl of a 10 M NaOH stock at 37 °C for 15 min. Samples were neutralized with 0.7 μl of a 7.5 M HCl stock and the remaining complementary DNA was purified using a nucleotide removal kit (QIAGEN). Sequencing reactions were performed using a commercial kit designed to be used with fluorescent dye-labeled primers (Thermo Fisher Scientific). ExoSAP-IT reagent (Thermo Fisher Scientific) was used to remove excess dNTPs and primers from the PCR product prior to the sequencing reaction. For the sequencing procedure, 4 μl of purified PCR product (or approximatively 0.5 – 1 pmol of DNA) and 2 pmol of the 5’-labeled oligonucleotide were used to prepare a master reaction following the manufacturer’s instructions. The following thermocycler program was used for the sequencing reaction: 30 s denaturation at 95°C; 15 s annealing at 50°C; 25 cycles of 60 s for elongation at 72°C. 2 μL of formamide loading dye from the kit were then added to each Sanger reaction, samples were heated for 3 min at 75°C to denature the cDNA, while the toeprinting samples were denatured at 95 °C for 5 min. 3.5 μl of the sequencing reactions and 3 μl of the toeprinting reactions were separated by 7.5% sequencing PAGE (2,000 V, 40 W for 2–2.5 h) followed by detection on an Amersham Typhoon Gel and Blot Imaging System (GE Healthcare Life Sciences).

### β-galactosidase complementation assay

To test for *in vivo* activity, a pERMZα^16^ plasmid containing the *ermCL-ermC* operon in frame with the lacZα reporter was used. A sequence coding for the MAAAPQKC peptide was inserted in place of the first ten codons of the ermCL sequence using the QuikChange Lightning Site-Directed Mutagenesis Kit (Agilent Technologies, Inc). Briefly, oligonucleotides (*pZα_MAAAPQKC_fwd*) and (*pZα_MAAAPQKC_rvs*) were used to mutate the pERMZα plasmid according to the manufacturer’s instructions. Both the pERMZα and pERMZα-MAAAPQKC plasmids were transformed into *E. coli* JM109 from which the *acrAB* operon had been deleted to reduce drug efflux^38,39^. Cells were grown in lysogeny broth (LB) at 37ºC (180 rpm) with ampicillin (100 μg/ml) until they reached an optical density of 0.5 at 600 nm. 2 ml of the cell culture were added to 8 ml of 0.6% LB-agar at 50ºC and plated onto 1.5% LB agar plates (120 mm diameter) supplemented with 0.5 mM ampicillin, 0.5 mM Isopropyl-β-D-1-thiogalactopyranoside (IPTG) and 0.5 mM 5-bromo-4-chloro-3-indolyl-beta D-galactopyranoside (X-gal). Once the soft agar had solidified, 6-mm-diameter Whatman paper discs (GE Healthcare) were placed on top of the plate and wetted with 5 μl of a 10 mM TcmX solution in DMSO or 300 μg of erythromycin diluted in 10 μl of ethanol. The plates were then incubated at 30°C overnight and pictures were taken the next day.

### Preparation of an *E. coli* 70S-MAAAPQKC-TcmX complex for cryo-EM

Stalled ribosomal complexes for cryo-EM were obtained by translating the PQKC-containing template used for toeprinting *in vitro*, in a reaction containing 2.2 μM of reassociated *E. coli* 70S ribosomes, 100 μM TcmX, 5 pmol of *TP_MAAAPQKCAAA\** mRNA (see “Toeprinting assays” section) and components of the PURExpress **Δ**Ribosomes Kit (New England Biolabs). The reaction was incubated at 37ºC for 20 min and then diluted in 50 mM Hepes KOH pH 7.5, 100 mM K-Acetate, 25 mM Mg-Acetate, 10 μM TcmX to yield a final ribosome concentration of 300 nM. Quantifoil carbon grids (QF-R2/2-Cu) were coated with a thin carbon layer (∼2 nm) using a Safematic ccu-010 HV carbon coater. Grids were glow-discharged for 30 s at 2 mA before application of 3.5 μl of 70S-MAAAPQKC-TcmX complex. After blotting for 2.5 s and waiting for 30 s (blotting force 5), grids were plunge-frozen in liquid ethane using an FEI Vitrobot Mark IV (Thermo Fisher Scientific) set to 4 °C and 100% humidity.

### Cryo-EM data acquisition

Cryo-EM images were collected in counting mode on a Talos Arctica (Thermo Fisher Scientific) operated at 200 kV and equipped with a K2 Summit direct electron detector (Gatan) in Nanoprobe mode at the IECB in Bordeaux (France). Images were recorded with SerialEM with a magnified pixel size of 0.93 Å at a magnification of 45,000x to record 38 movie frames with an exposure time of 3.8 seconds using a dose rate of 0.94 e^−^/Å^2^ per frame for a total accumulated dose of 35.72 e^−^/Å^2^ (Supplementary Table 2). The final dataset was composed of 7,450 micrographs with defocus values ranging from -0.4 to -2.5 μm.

### Cryo-EM image processing

Data were processed in Relion v4.0^40^ according to the scheme presented in Extended Data Fig. 6. Briefly, the raw movie frames were summed and corrected for drift and beam-induced motion at the micrograph level using the Relion v4.0^40^ implementation of MotionCor2^41^. The resolution range of each micrograph and the contrast transfer function (CTF) were estimated with CTFFIND v4.1^42^. Well-defined two-dimensional classes were selected by subsequent rounds of two-dimensional classification of the particles obtained by automated picking in Relion v4.0^40^. Three-dimensional classification was performed in Relion v4.0^40^ in three steps: (1) unsupervised classification with particles downsized four times; (2) focused classification with background subtraction using a mask covering all three tRNA sites and the GTPase binding site on particles downsized twice; and (3) focused classification with background subtraction using a mask covering the A site on particles downsized twice. Classes containing tRNAs in the A, P and E sites were selected for automated 3D reconstruction and CTF refinement in Relion v4.0^40^, followed by Bayesian polishing. The final reconstruction was sharpened by applying a negative sharpening B-factor of -10 in Relion v4.0^40^. The resolution for the electron density map was estimated using the gold standard criterion (FSC=0.143) resulting in a final reconstruction with a nominal resolution of 2.7 Å. Local-resolution estimation was performed using Relion v4.0^40^. The pixel size was optimized by calculating model-to-map correlation coefficients at multiple pixel sizes in Chimera v1.16^43^, and selecting the pixel size that yielded in the highest coefficient value.

### Atomic model building and refinement

An initial model of the 70S-TcmX complex was obtained by placing the coordinates of an *E. coli* 70S ribosome (PDB: 6TBV) and of the TcmX antibiotic (PDB: 6Y69) into an auto-sharpened map obtained with Phenix v1.20.1^44^. The nascent chain, mRNA and A-, P- and E-site tRNAs were modeled *de novo* into the corresponding density using Coot v0.9.7^45^. The model was refined using Isolde v1.3^46^ and the real space refinement procedure in Phenix v1.20.1^47^, with reference model restraints.

### Figure preparation

Plots were produced with Python3 and Matplotlib 3.3.2^36^. An auto-sharpened map from Phenix v1.20.1^44^ was used to prepare all figures showing cryo-EM density except Extended Data Fig. 6 and 7, for which maps obtained directly from Relion v4.0^40^ were used. Figures showing cryo-EM density or atomic models were prepared using ChimeraX v.1.3^48^ or PyMOL v.2.5.0 (Schrödinger).

## Supporting information

Source Data for Fig. 1d

Source Data for Fig. 2a

Source Data for Fig. 2b

## ACKNOWLEDGEMENTS

We thank Ilya Osterman and Daniel Wilson for providing TcmX, Nora Vázquez-Laslop for providing the JM109 Δ*acrA*/*acrB E. coli* strain, Alok Malhotra for providing RNase R, and Scott Blanchard for his comments on an early draft of the manuscript. We also thank Anaïs Labécot for preliminary experiments. E.C.L. and C.A.I. have received funding for this project from the European Research Council (ERC) under the European Union’s Horizon 2020 research and innovation program (Grant Agreement No. 724040). C.A.I. is an EMBO YIP and has received funding from the Fondation Bettencourt-Schueller. T.N.P. has received funding for this project from the Agence Nationale de la Recherche (ANR) under the frame of the Joint JPI-EC-AMR Project “Ribotarget – Development of novel ribosome-targeting antibiotics”. We thank the cryo-EM facility of the European Institute for Chemistry and Biology (Pessac, France) for the data collection time on the Talos Arctica microscope.

## AUTHOR CONTRIBUTIONS

C.A.I. designed the study. E.C.L. performed iTP-seq and toeprinting experiments. E.C.L. and T.T.R. processed and analyzed the iTP-seq data. T.N.P. and T.T.R. performed the blue ring assays. E.C.L. prepared complexes for cryo-EM; T.N.P. prepared grids and performed cryo-EM data collection. T.N.P. and C.A.I. processed the cryo-EM data, and E.C.L. and C.A.I. built the model. E.C.L., T.N.P., T.T.R. and C.A.I. wrote the manuscript.

## COMPETING FINANCIAL INTERESTS

The authors declare no competing financial interests. Correspondence and requests for materials should be addressed to T.T.R. or C.A.I.

## DATA AVAILABILITY

Cryo-EM maps of the 70S-MAAAPQKC-TcmX complex and the associated molecular models have been deposited in the Electron Microscopy Data Bank and Protein Data Bank, with the accession codes EMD-14956 and PDB 7ZTA, respectively. Sequencing data for the iTP-seq experiment have been deposited in the National Center for Biotechnology Information Short Read Archive with the accession code PRJNA854319. Source data are provided with this paper.

## CODE AVAILABILITY

Scripts used to process iTP-seq data were deposited on GitHub (https://github.com/innislab/ITP-seq).

**Extended Data Fig. 1.**
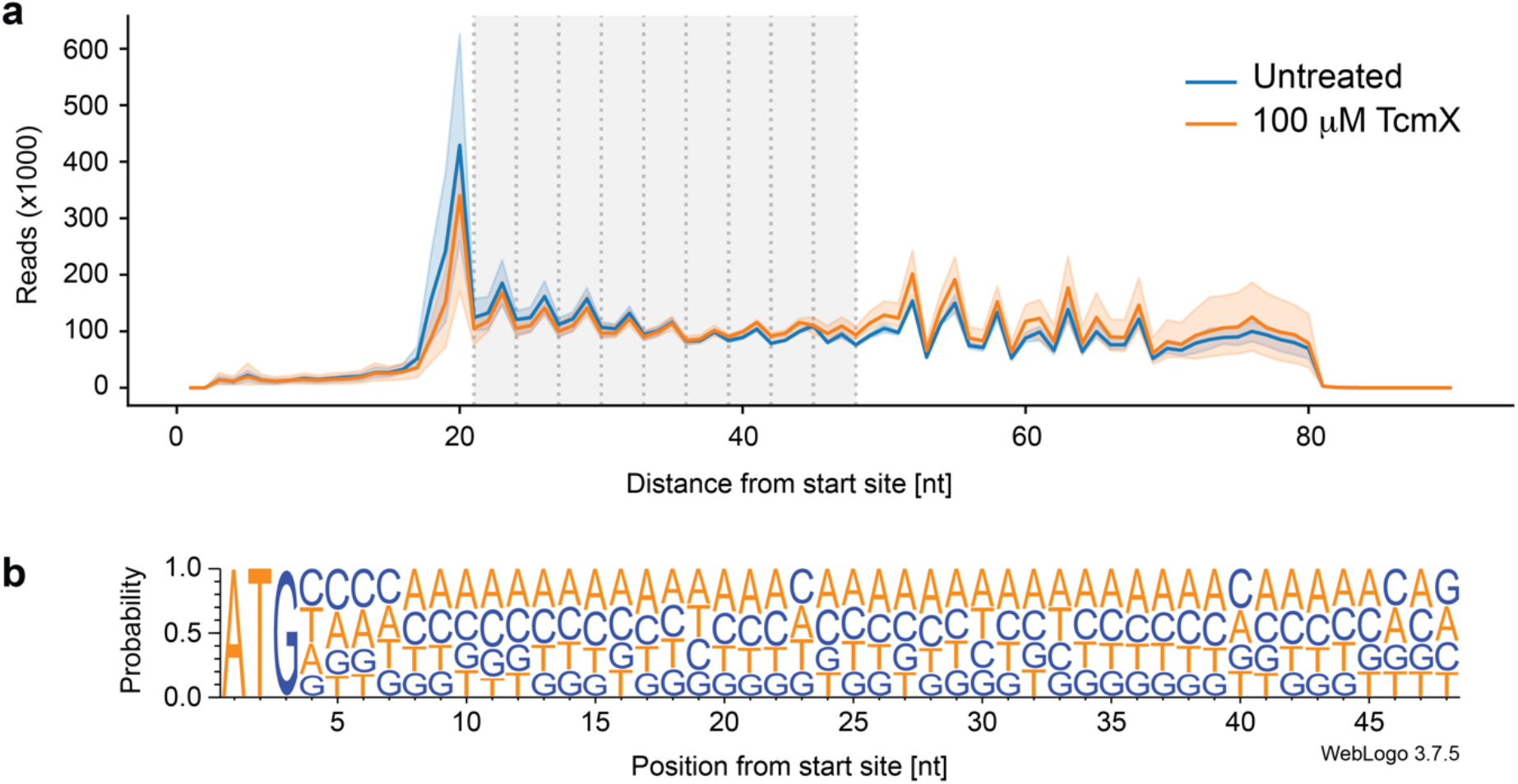
Size distribution of inverse toeprints and diversity of the (NNN)_15_ library. **a**, Number of deep sequencing reads obtained for inverse toeprints of different lengths, plotted as a function of the distance between the start codon and the RNase R cleavage site. Average read numbers for three independent replicates are shown for TcmX-treated (orange) and untreated (blue) samples. Inverse toeprints within a size range featuring a well-defined three-nucleotide periodicity were retained for further analysis (gray area). The iTP-seq experiment consisted of 3 independent replicates for the TcmX-treated condition (2.8×10^6^, 2.5×10^6^ and 2.3×10^6^ reads), and 3 replicates for the untreated condition (2.7×10^6^, 2.3×10^6^ and 2.6×10^6^ reads). On average ∼2.5×10^6^ reads were obtained per replicate. The error bands show the 95% confidence interval of the mean of the three replicates. **b**. Nucleotide frequency of the raw reads for the (NNN)_15_ library, showing a relatively equal proportion of each dNTP for codons 2-15.

**Extended Data Fig. 2.**
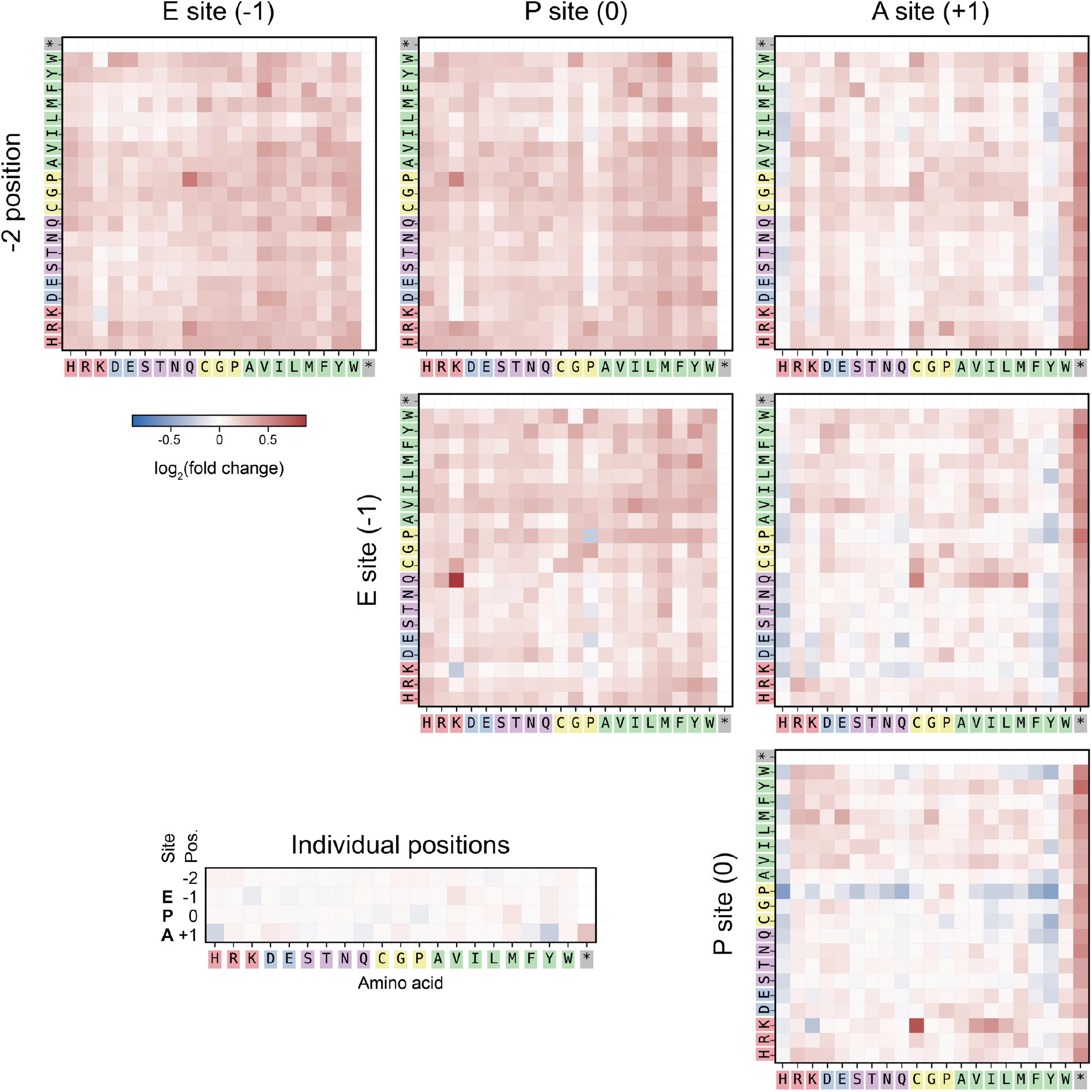
Enrichment of two-amino acid motifs in samples treated with TcmX. Heatmaps showing the enrichment of individual residues at positions -2 to +1, and all possible combinations of motifs consisting of two adjacent amino acids, following treatment with TcmX. Enrichment is defined as log_2_(F_TcmX_/F_untreated_), where F_TcmX_ is the frequency of occurrence of a given motif in the sample treated with TcmX and F_untreated_ is its frequency in the untreated sample.

**Extended Data Fig. 3.**
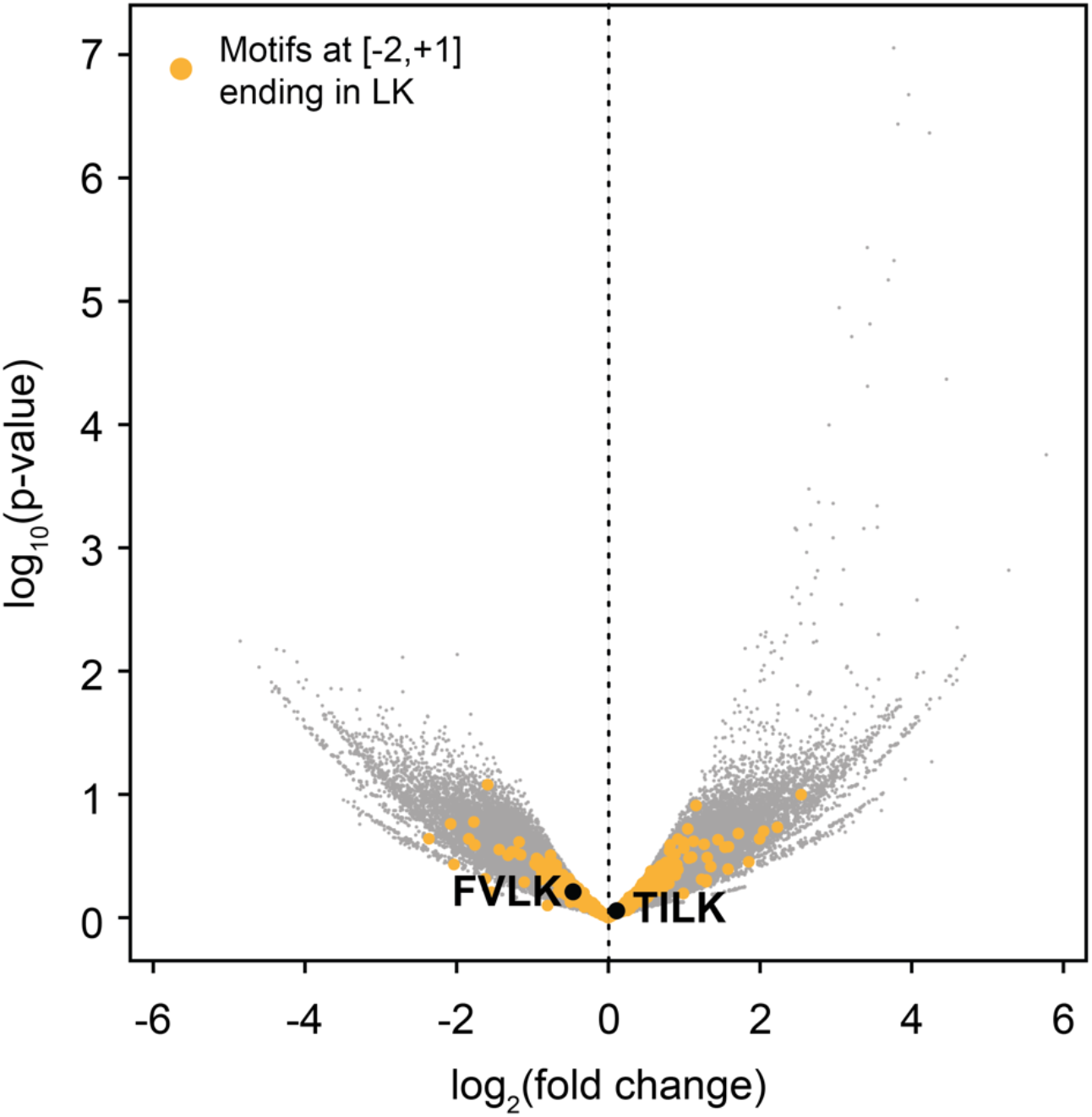
Four-amino acid motifs at position [-2,+1] ending with LK are not significantly enriched following TcmX treatment. Volcano plot of statistical significance (-log_10_(p-value)) against enrichment (log_2_(fold change)) for four-amino acid motifs at position [-2,+1]. Motifs ending in LK are shown in yellow and the two LK-containing motifs described in Ref. 6 (FVLK/TILK) are labeled in bold.

**Extended Data Fig. 4.**
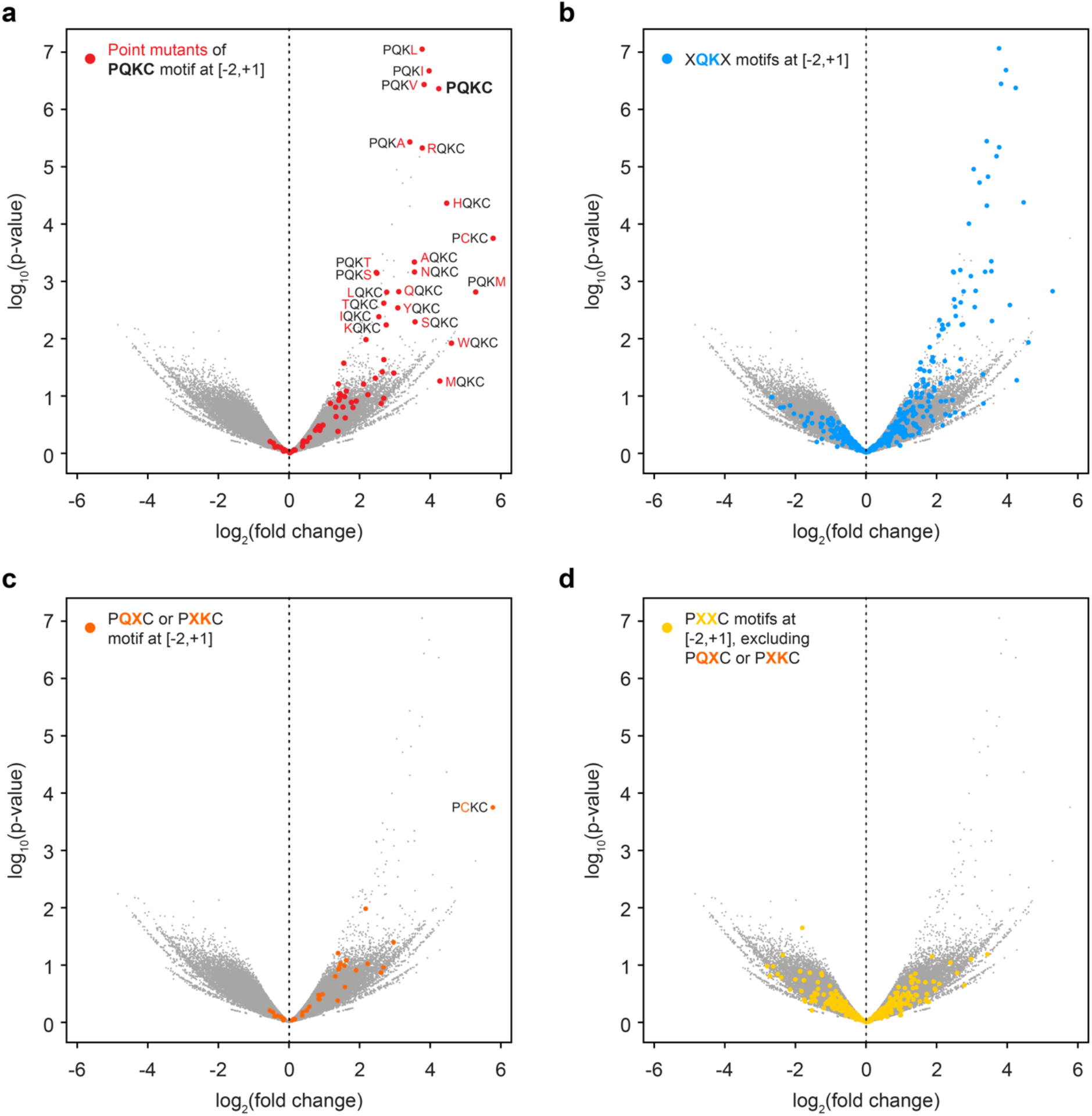
Ability of different PQKC variants to induce TcmX-dependent stalling. Volcano plots of statistical significance (-log_10_(p-value)) against enrichment (log_2_(fold change)) for four-amino acid motifs at position [-2,+1]. Single amino acid variants of the PQKC motif are shown in red, XQKX motifs are in blue, PQXC or PXKC motifs are in orange and PXXC motifs are in yellow.

**Extended Data Fig. 5.**
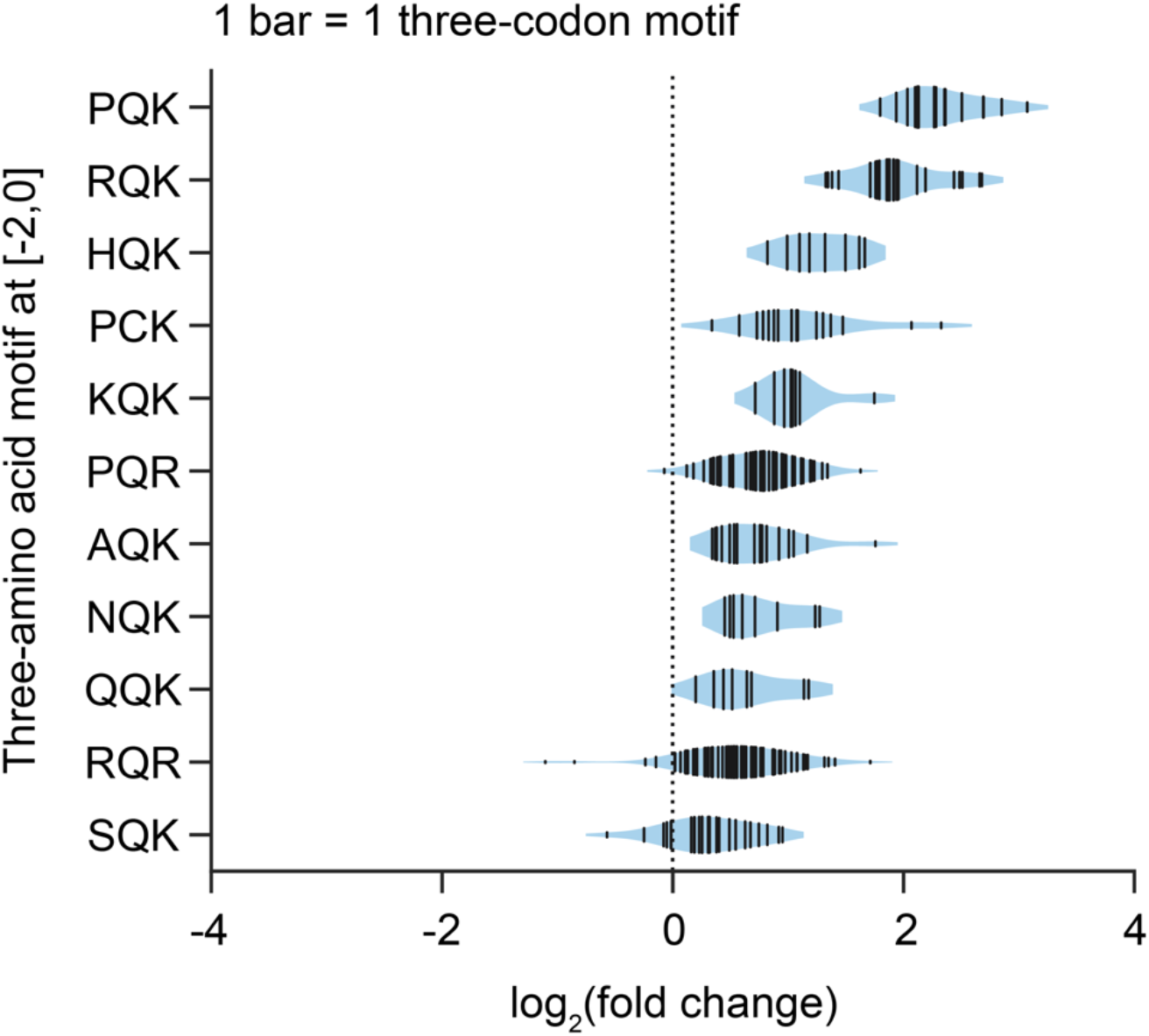
Codon composition does not have a major influence on TcmX-dependent ribosome stalling at QK motifs. Violin plot showing the enrichment of various three-amino acid motifs at positions [-2,0] following treatment with TcmX. Each bar represents a motif variant with a distinct codon composition.

**Extended Data Fig. 6.**
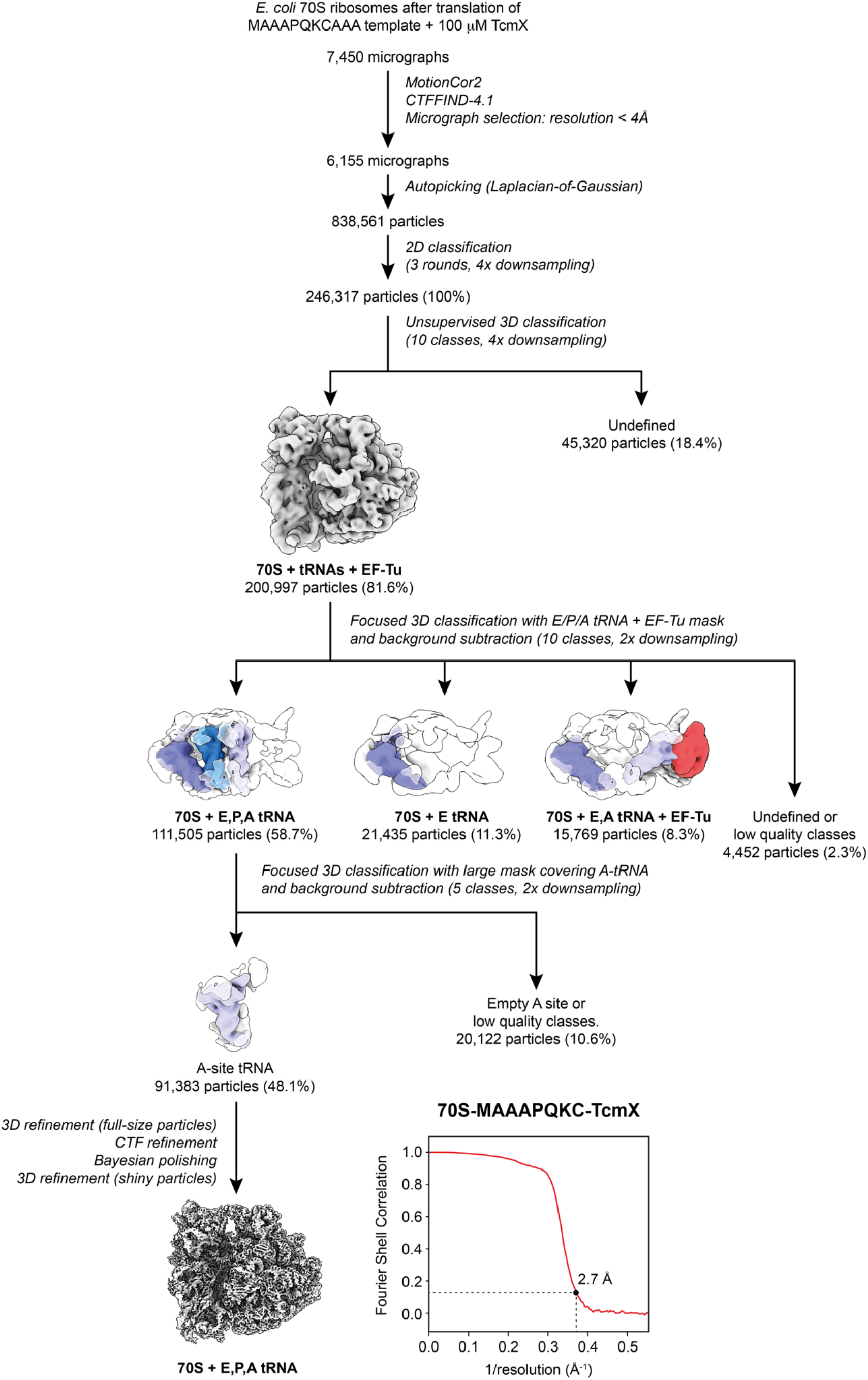
Overview of cryo-EM data processing. Flowchart showing the workflow used to process cryo-EM data for the 70S-MAAAPQKC-TcmX complex in Relion v4.0 (Ref. 42). The final reconstruction could be refined to an overall resolution of 2.7 Å, assessed using a Fourier shell correlation (FSC) cutoff of 0.143.

**Extended Data Fig. 7.**
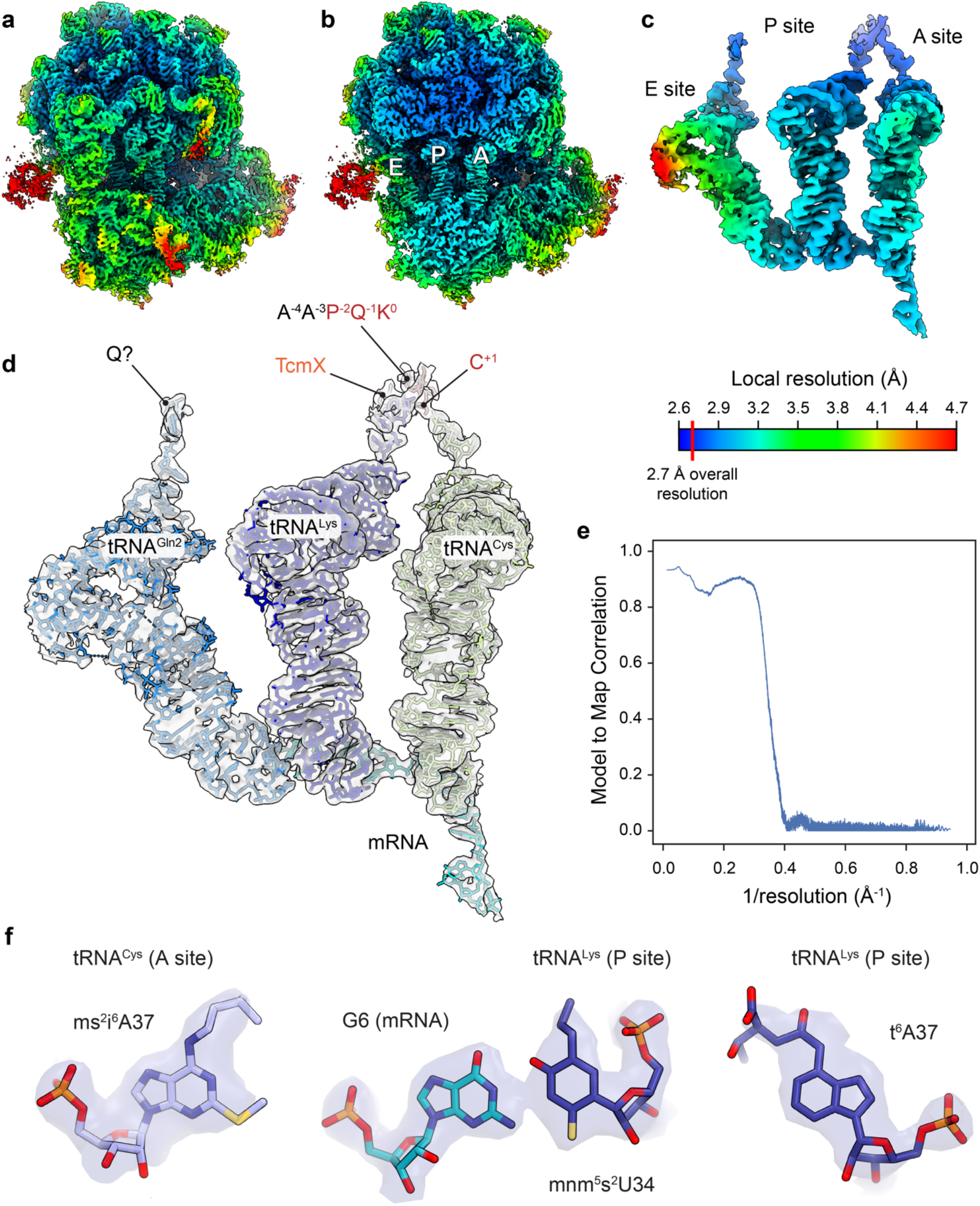
Quality of the 70S-MAAAPQKC-TcmX reconstruction. **a**, Cryo-EM density map obtained for a 3D reconstruction of the 70S-MAAAPQKC-TcmX complex in Relion v4.0 (Ref. 42) and auto-sharpened with Phenix v1.20.1 (Ref. 46), filtered and colored by local resolution estimation values in Chimera X v.1.3 (Ref. 50). **b**, Cross-section of the same map showing the E, P and A sites of the ribosome. **c**, Isolated ligand densities in the E, P and A sites, filtered and colored by local resolution. **d**, The same isolated densities shown in c, displayed as a transparent surface and fitted with molecular models of the E-, P- and A-site tRNAs, mRNA, TcmX and the nascent peptide. **e**, Model to map correlation curve calculated for the 70S-MAAAPQKC-TcmX structure. **f**, Representative cryo-EM densities of three different tRNA modifications within the A- and P-site tRNAs.

**Extended Data Fig. 8.**
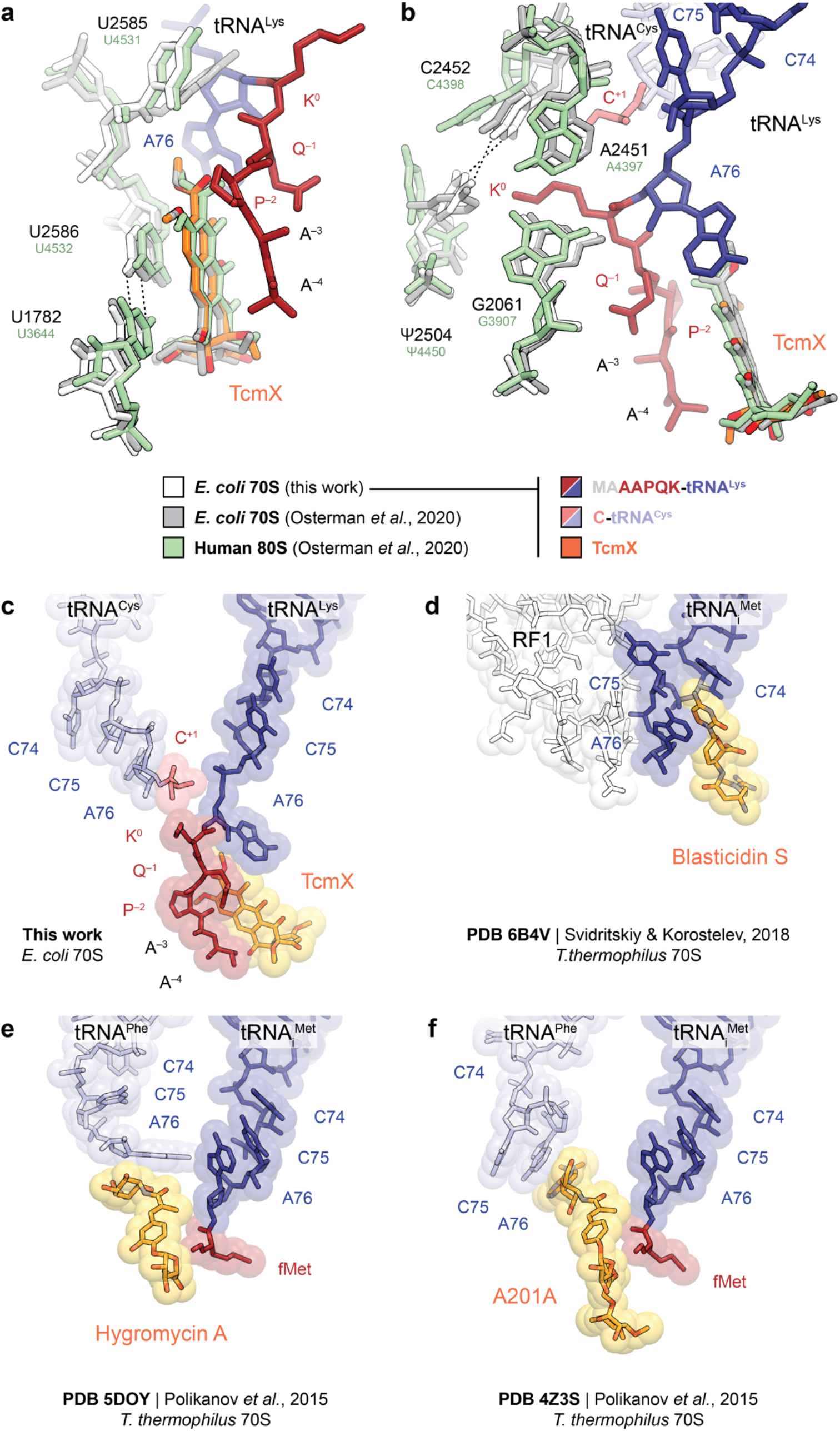
Comparison of 70S-MAAAPQKC-TcmX with earlier structures of the ribosome in complex with TcmX or other antibiotics. **a**,**b**, Structural comparison of the peptidyl transferase center and TcmX binding site in 70S-MAAAPQKC-TcmX (white) and in complexes between *E. coli* 70S (gray, PBD 6Y69) or human 80S (pale green, PDB 6Y6X) ribosomes and TcmX (Ref. 6). **c**,**d**, Displacement of the 3’ end of peptidyl-tRNA (dark blue) by TcmX (c) or Blasticidin S (PDB 6B4V, Ref. 23). **e**,**f**, Displacement of the 3’ end of aminoacyl-tRNA (light blue) by (e) Hygromycin A (PDB 5DOY, Ref. 25) or (f) A201A (PDB 4Z3S, Ref. 25).

**Extended Data Fig. 9.**
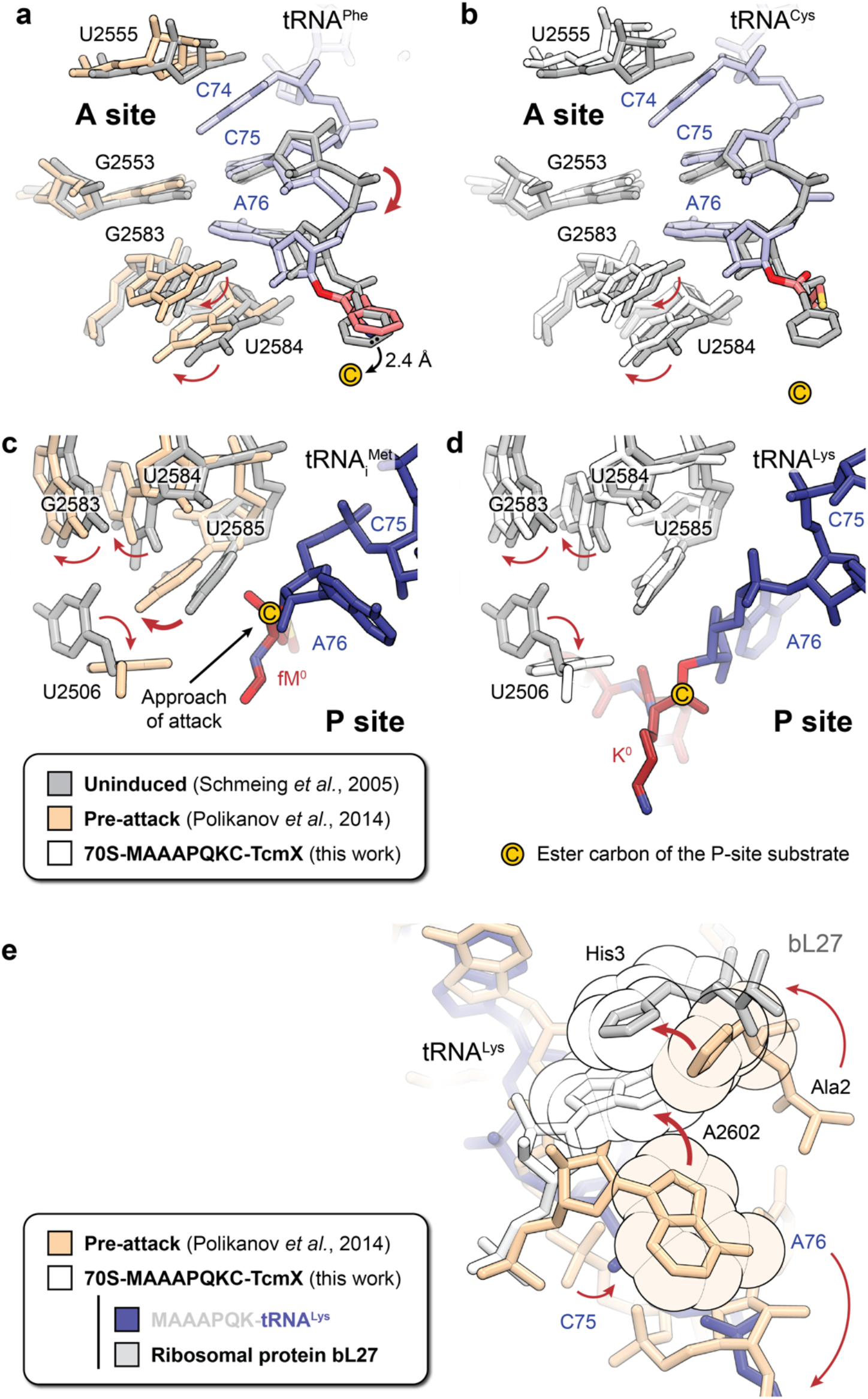
Effect of TcmX on the accommodation of the 3’ end of aminoacyl-tRNA, on 23S rRNA residue 2602 and on the N-terminus of ribosomal protein bL27. **a-d**, Structural comparison of 23S rRNA nucleotides and tRNAs within the A site (a,b) and P site (c,d) regions of the peptidyl transferase center in the uninduced (Ref. 21) and induced (pre-attack) (Ref. 22) states (PDB 1VQ6 and 1VY4, respectively) (a,c), and the uninduced and TcmX-inhibited states (b,d). In the 70S-MAAAPQKC-TcmX structure, 23S rRNA residues U2506, G2583 and U2584 are in their induced conformation, but 23S rRNA nucleotide U2585 and nucleotide A76 of the P-site tRNA remain in the uninduced conformation, indicating partial aminoacyl-tRNA accommodation into the peptidyl transferase center. **e**, Sequestration of residue A76 of the P-site tRNA^Lys^ within the tunnel by TcmX causes the preceding residue C75 to shift, which in turn leads to a conformational change in the peptidyl transferase center resulting in the stacking of the base of 23S rRNA residue A2602 against the imidazole ring of residue His-3 of ribosomal protein bL27.

**Extended Data Fig. 10.**
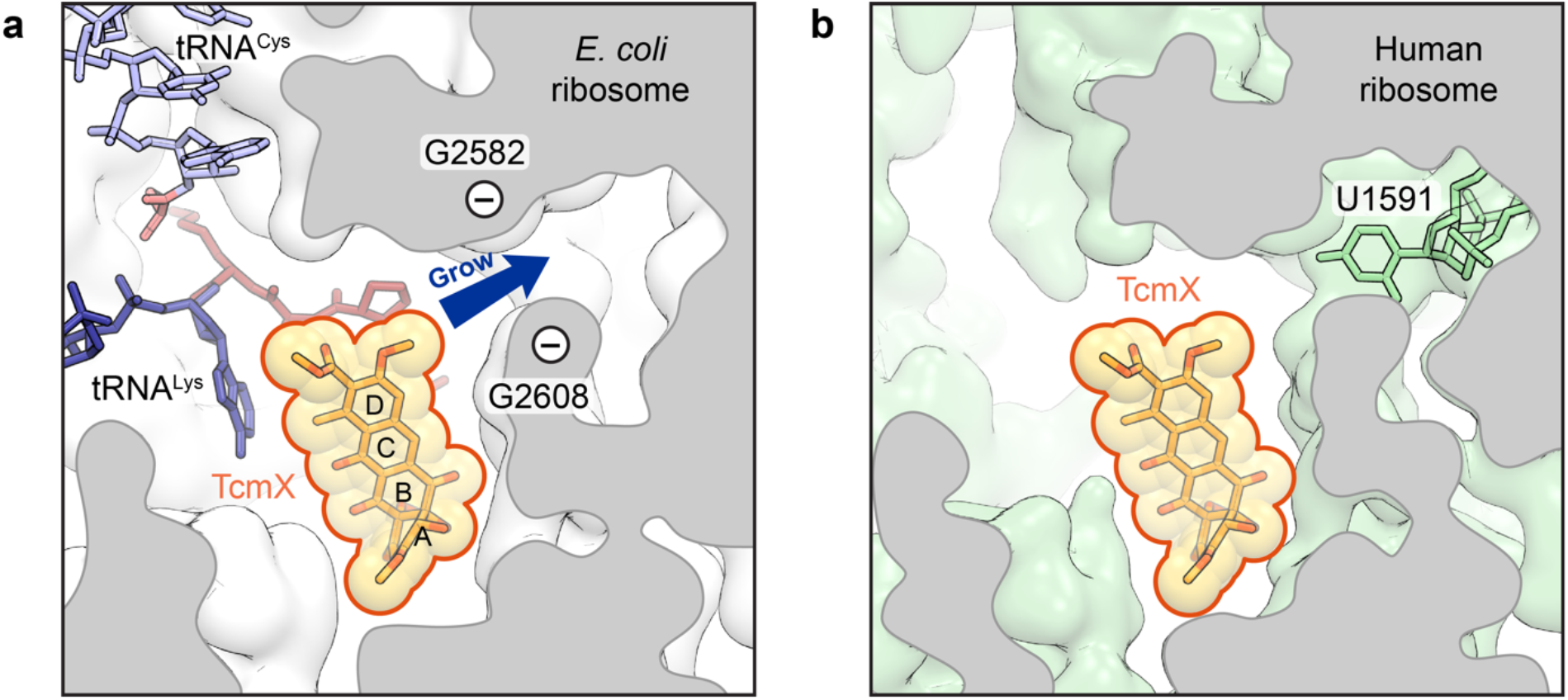
Possible way forward for the design of TcmX derivatives with increased specificity for the bacterial ribosome. **a**, The O-methyl group attached to the D ring of TcmX could be used as a starting point to “grow” the drug molecule towards a cavity lined with the backbone phosphate groups of 23S rRNA residues G2582 and G2608 in the *E. coli* ribosome (white). **b**, TcmX derivatives with suitable side chains attached to this position may retain activity against the bacterial ribosome while no longer binding to the human ribosome (pale green, PDB 6Y6X), where this cavity is blocked by 28S rRNA residue U1591.

## SUPPLEMENTARY INFORMATION

**Supplementary Table 1.**
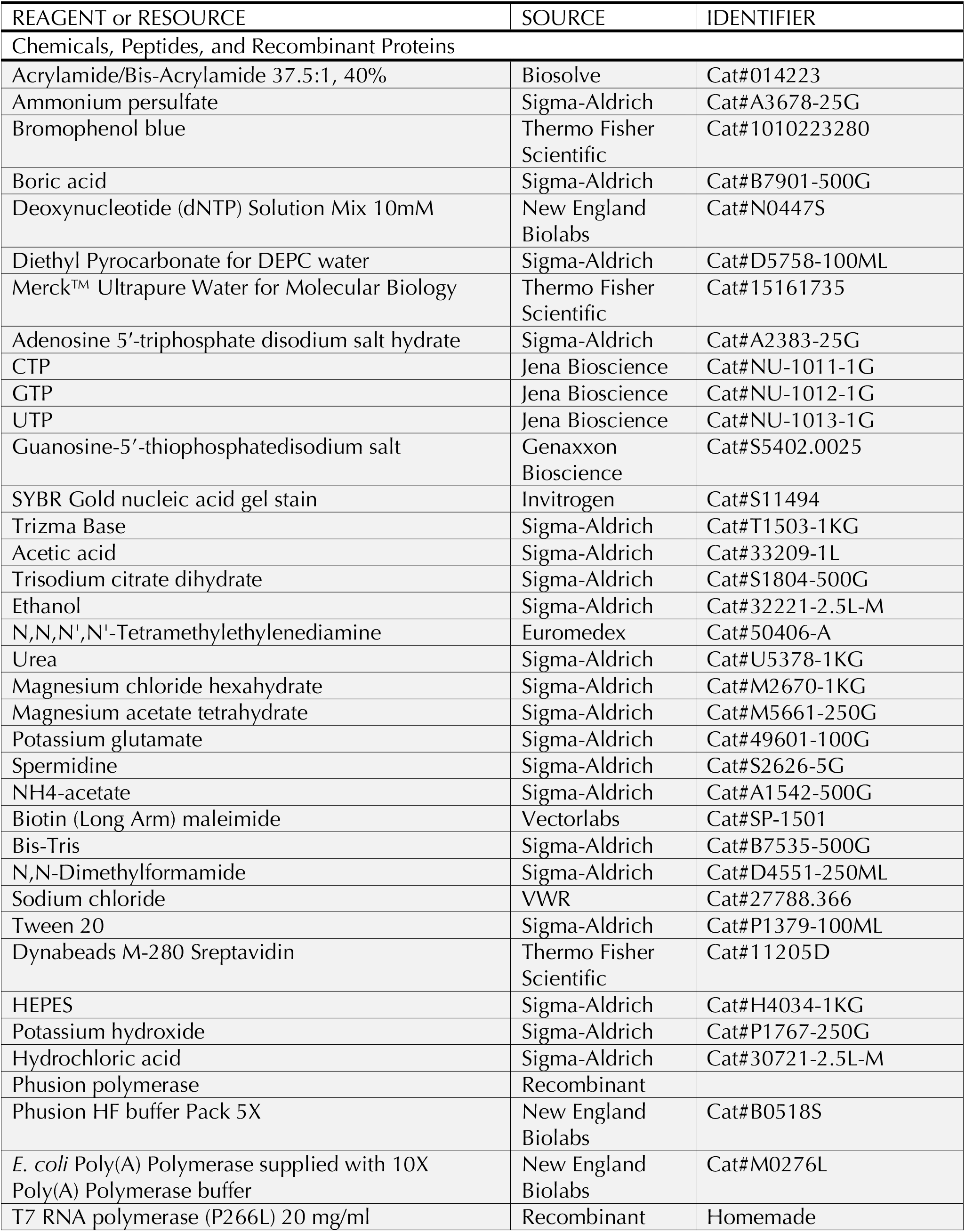
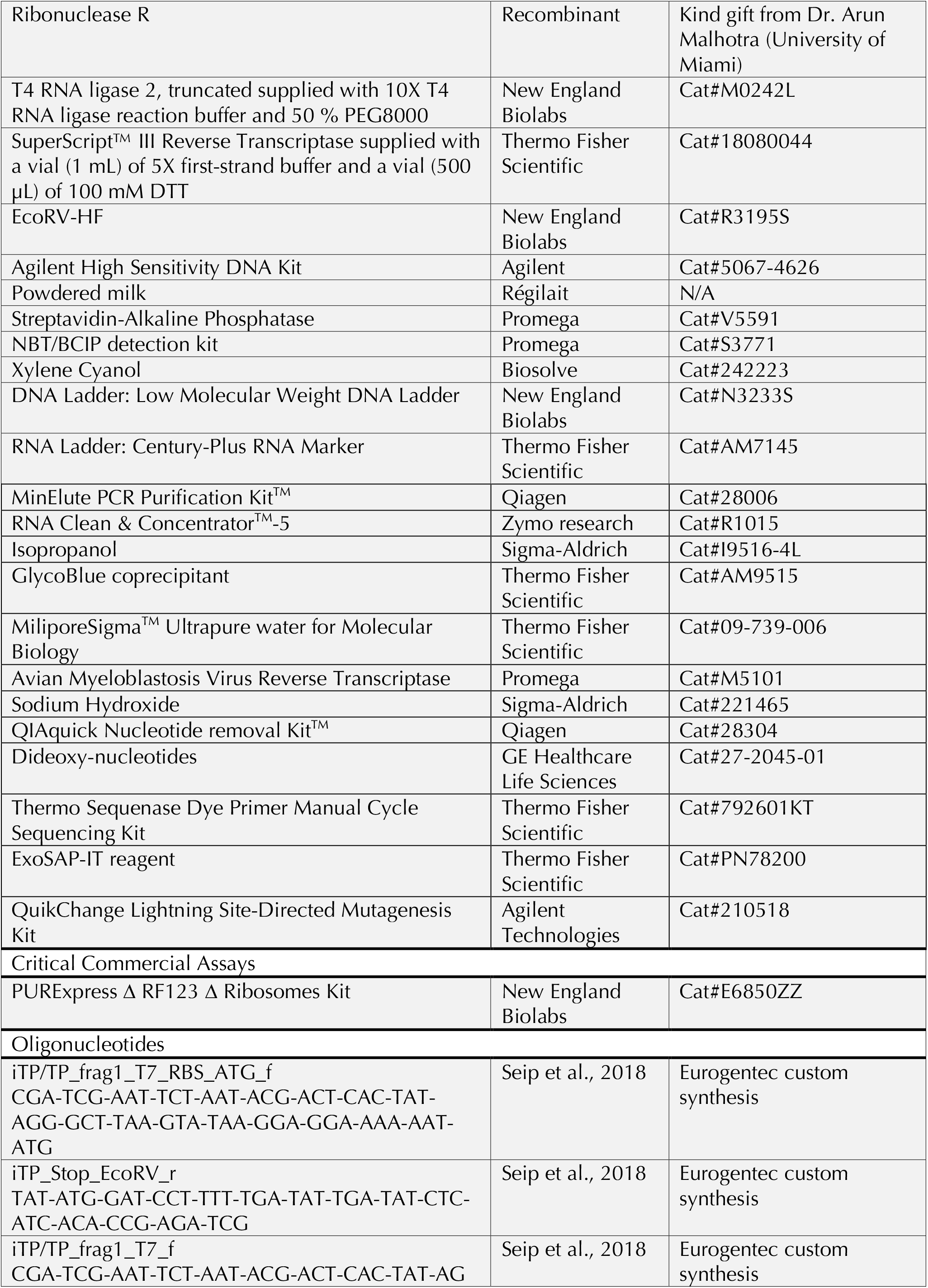
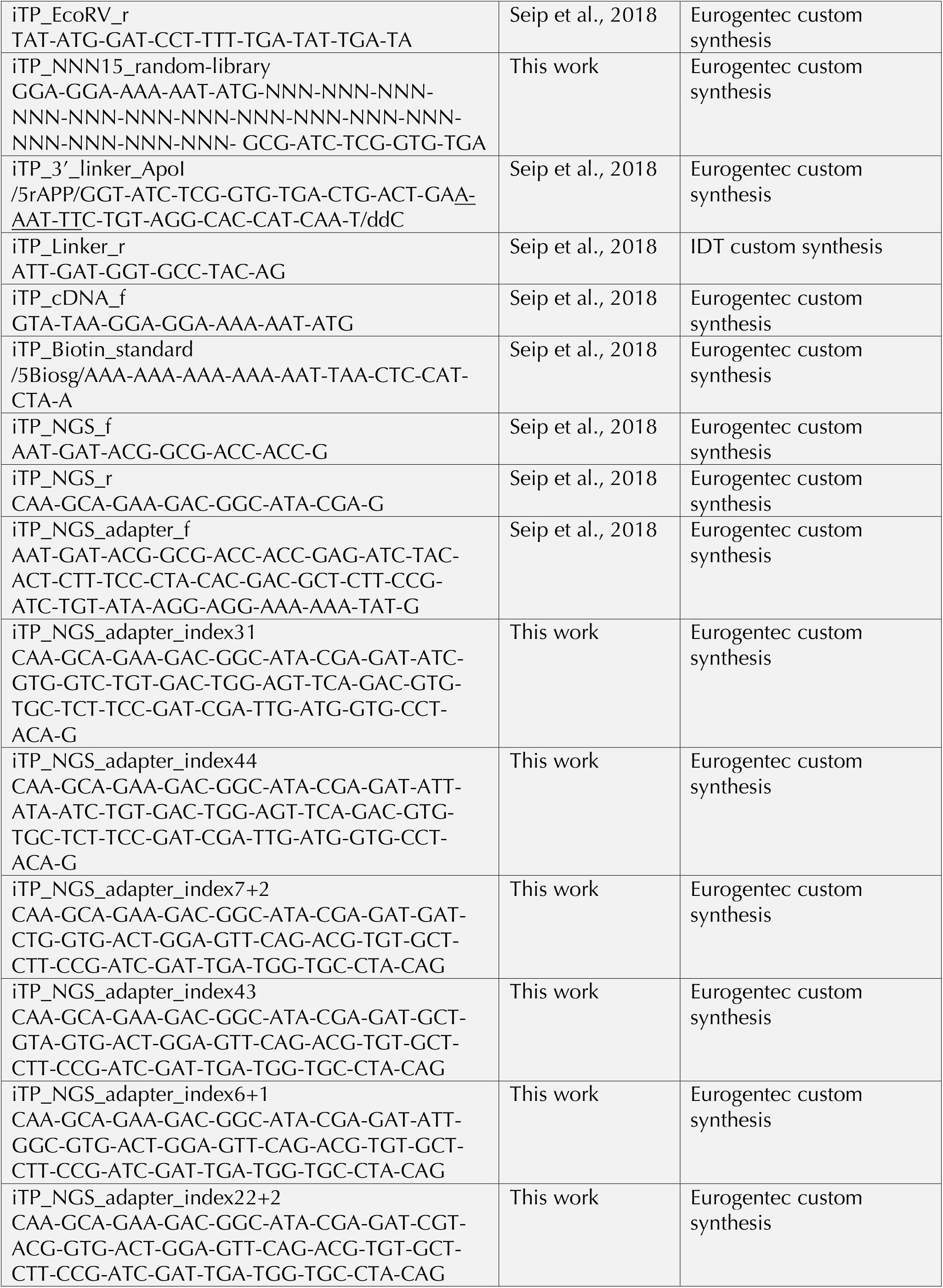
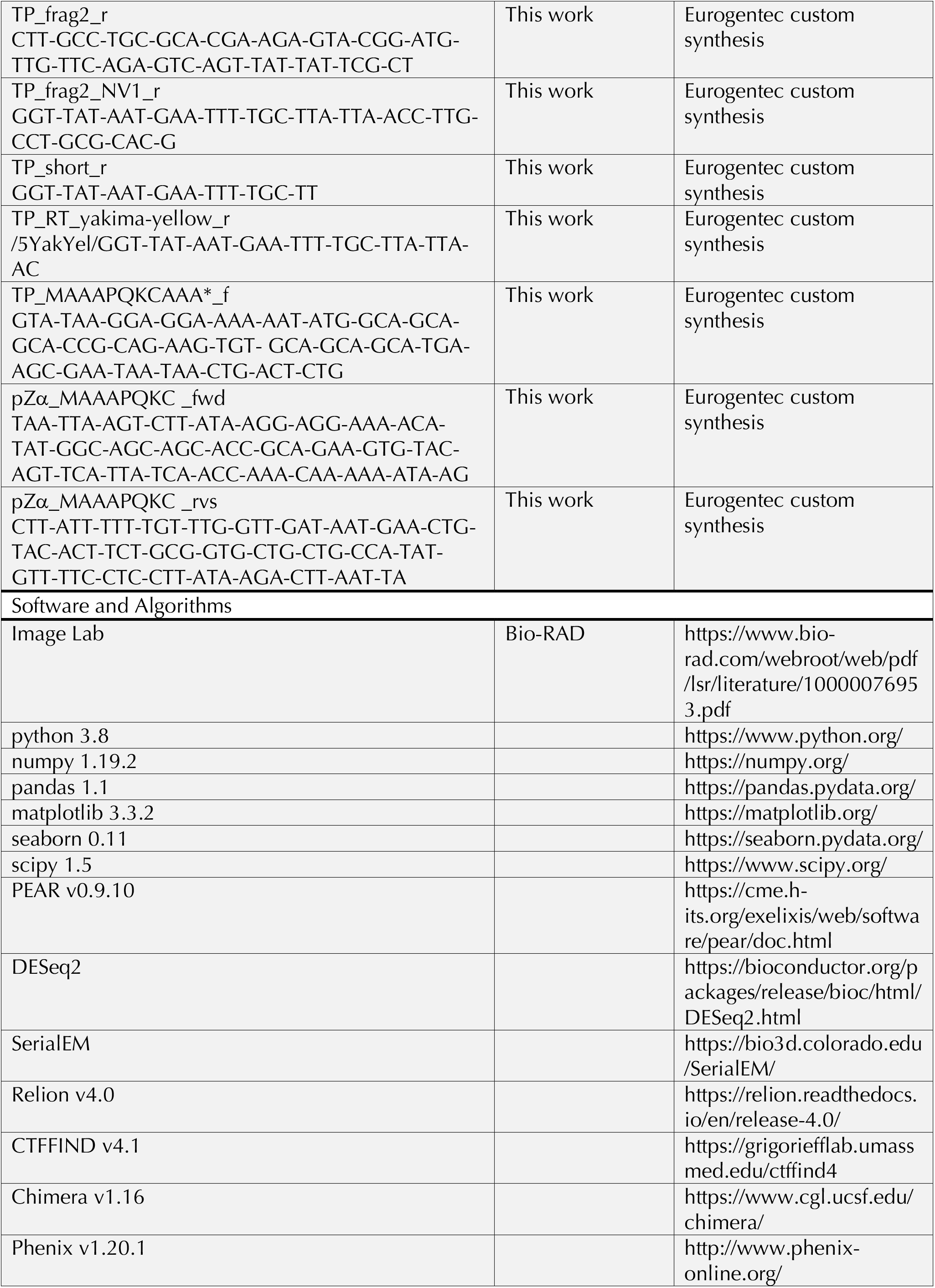
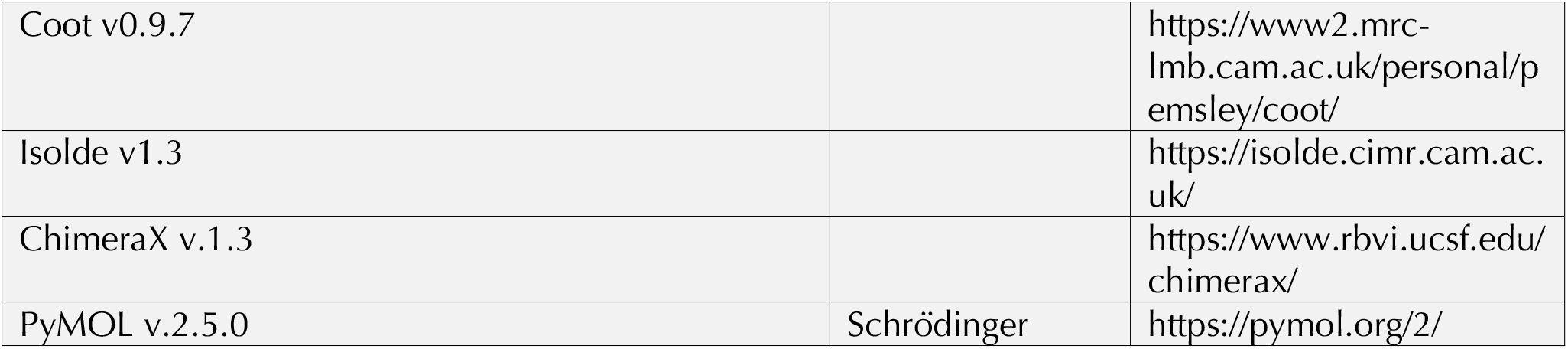
Reagents, oligonucleotides and software.

**Supplementary Table 2.**
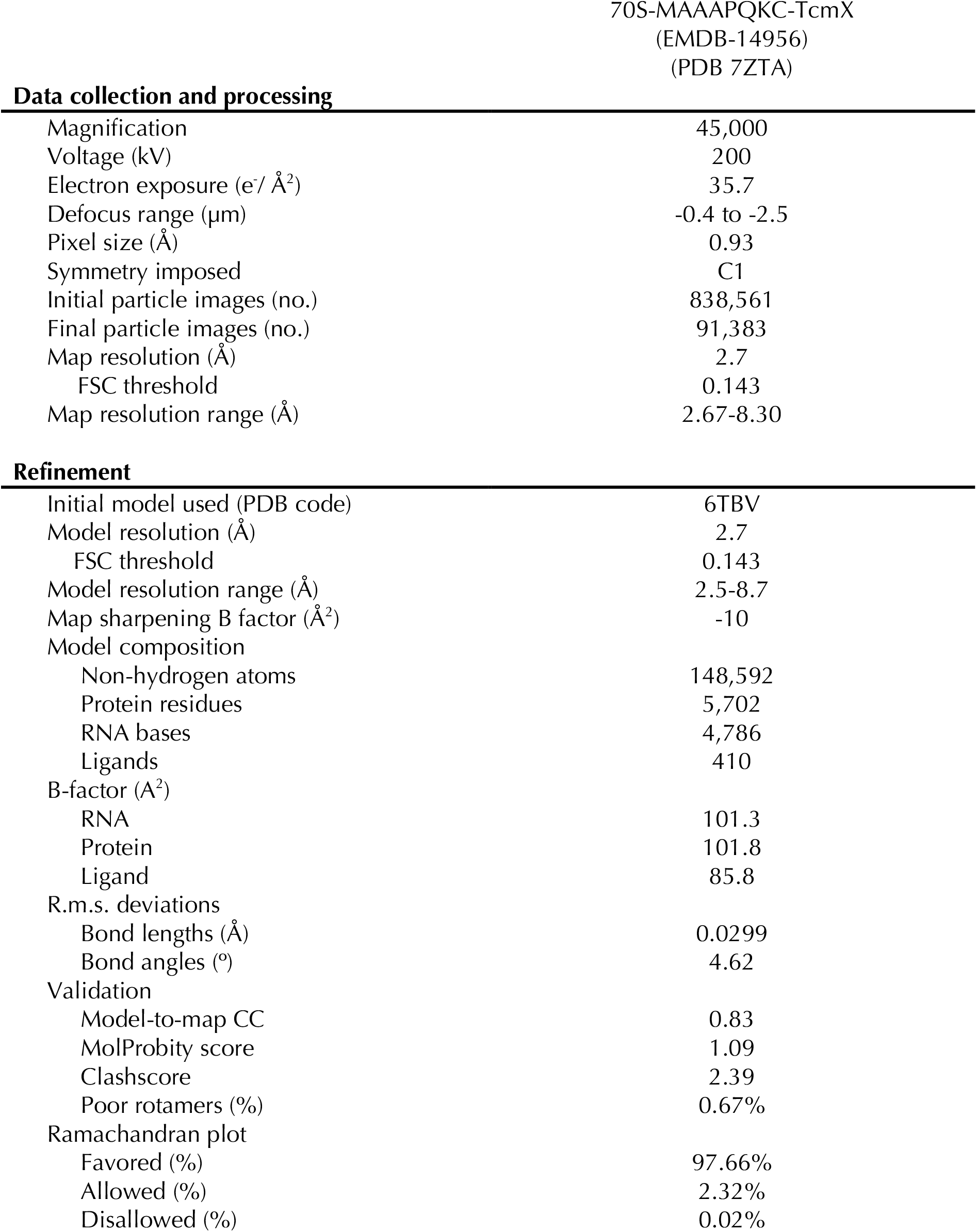
Cryo-EM statistics.

Data collection and processing
|  |  |
| --- | --- |
| Magnification | 45,000 |
| Voltage (kV) | 200 |
| Electron exposure (e<sup>-</sup>/ Å<sup>2</sup>) | 35.7 |
| Defocus range (μm) | -0.4 to -2.5 |
| Pixel size (Å) | 0.93 |
| Symmetry imposed | C1 |
| Initial particle images (no.) | 838,561 |
| Final particle images (no.) | 91,383 |
| Map resolution (Å) | 2.7 |
| FSC threshold | 0.143 |
| Map resolution range (Å) | 2.67-8.30 |
| Refinement |  |
| Initial model used (PDB code) | 6TBV |
| Model resolution (Å) | 2.7 |
| FSC threshold | 0.143 |
| Model resolution range (Å) | 2.5-8.7 |
| Map sharpening B factor (Å<sup>2</sup>) | -10 |
| Model composition |  |
| Non-hydrogen atoms | 148,592 |
| Protein residues | 5,702 |
| RNA bases | 4,786 |
| Ligands | 410 |
| B-factor (Å<sup>2</sup>) |  |
| RNA | 101.3 |
| Protein | 101.8 |
| Ligand | 85.8 |
| R.m.s. deviations |  |
| Bond lengths (Å) | 0.0299 |
| Bond angles (°) | 4.62 |
| Validation |  |
| Model-to-map CC | 0.83 |
| MolProbity score | 1.09 |
| Clashscore | 2.39 |
| Poor rotamers (%) | 0.67% |
| Ramachandran plot |  |
| Favored (%) | 97.66% |
| Allowed (%) | 2.32% |
| Disallowed (%) | 0.02% |

## Notes

### Competing Interest Statement

The authors have declared no competing interest.

### Summary of Updates

Additional text and extended data figures were included to show the effect of various mutations to the PQKC motif on the ability of TcmX to stall translation (Extended Data Fig. 4), to provide additional structural detail (Extended Data Figs. 8 and 9) and to show a possible way forward for the development of TcmX derivatives with increased specificity for the bacterial ribosome. In addition, both the text and figures were edited to improve the overall clarity of the manuscript.

## REFERENCES

1. Huang, C. et al. Marine Bacterial Aromatic Polyketides From Host-Dependent Heterologous Expression and Fungal Mode of Cyclization. Front. Chem. 0, (2018).

2. Katz, L. & Baltz, R. H. Natural product discovery: past, present, and future. J. Ind. Microbiol. Biotechnol. 43, 155–176 (2016).

3. Wang, J. et al. Biosynthesis of aromatic polyketides in microorganisms using type II polyketide synthases. Microb. Cell Factories 19, 110 (2020).

4. Zhang, Z., Pan, H.-X. & Tang, G.-L. New insights into bacterial type II polyketide biosynthesis. F1000Research 6, 172 (2017).

5. Liu, B. et al. [Identification of tetracenomycin X from a marine-derived Saccharothrix sp. guided by genes sequence analysis]. Yao Xue Xue Bao 49, 230–236 (2014).

6. Osterman, I. A. et al. Tetracenomycin X inhibits translation by binding within the ribosomal exit tunnel. Nat. Chem. Biol. 16, 1071–1077 (2020).

7. Qiao, X. et al. Tetracenomycin X Exerts Antitumour Activity in Lung Cancer Cells through the Downregulation of Cyclin D1. Mar. Drugs 17, 63 (2019).

8. Jenner, L. et al. Structural basis for potent inhibitory activity of the antibiotic tigecycline during protein synthesis. Proc. Natl. Acad. Sci. 110, 3812–3816 (2013).

9. Kannan, K. et al. The general mode of translation inhibition by macrolide antibiotics. Proc. Natl. Acad. Sci. 111, 15958–15963 (2014).

10. Davis, A. R., Gohara, D. W. & Yap, M.-N. F. Sequence selectivity of macrolide-induced translational attenuation. Proc. Natl. Acad. Sci. U. S. A. 111, 15379–15384 (2014).

11. Seip, B., Sacheau, G., Dupuy, D. & Innis, C. A. Ribosomal stalling landscapes revealed by high-throughput inverse toeprinting of mRNA libraries. Life Sci. Alliance 1, e201800148 (2018).

12. Shimizu, Y. et al. Cell-free translation reconstituted with purified components. Nat. Biotechnol. 19, 751–755 (2001).

13. Starosta, A. L. et al. Translational stalling at polyproline stretches is modulated by the sequence context upstream of the stall site. Nucleic Acids Res. 42, 10711–10719 (2014).

14. Hartz, D., McPheeters, D. S., Traut, R. & Gold, L. Extension inhibition analysis of translation initiation complexes. Methods Enzymol. 164, 419–425 (1988).

15. Orelle, C. et al. Tools for Characterizing Bacterial Protein Synthesis Inhibitors. Antimicrob. Agents Chemother. 57, 5994–6004 (2013).

16. Bailey, M., Chettiath, T. & Mankin, A. S. Induction of erm(C) expression by noninducing antibiotics. Antimicrob. Agents Chemother. 52, 866–874 (2008).

17. Horinouchi, S. & Weisblum, B. Posttranscriptional modification of mRNA conformation: Mechanism that regulates erythromycin-induced resistance. Proc. Natl. Acad. Sci. 77, 7079– 7083 (1980).

18. Shivakumar, A. G., Hahn, J., Grandi, G., Kozlov, Y. & Dubnau, D. Posttranscriptional regulation of an erythromycin resistance protein specified by plasmic pE194. Proc. Natl. Acad. Sci. 77, 3903–3907 (1980).

19. Vazquez-Laslop, N., Thum, C. & Mankin, A. S. Molecular Mechanism of Drug-Dependent Ribosome Stalling. Mol. Cell 30, 190–202 (2008).

20. Gupta, P., Kannan, K., Mankin, A. S. & Vázquez-Laslop, N. Regulation of Gene Expression by Macrolide-Induced Ribosomal Frameshifting. Mol. Cell 52, 629–642 (2013).

21. Schmeing, T. M., Huang, K. S., Strobel, S. A. & Steitz, T. A. An induced-fit mechanism to promote peptide bond formation and exclude hydrolysis of peptidyl-tRNA. Nature 438, 520–524 (2005).

22. Polikanov, Y. S., Steitz, T. A. & Innis, C. A. A proton wire to couple aminoacyl-tRNA accommodation and peptide bond formation on the ribosome. Nat. Struct. Mol. Biol. 21, 787–793 (2014).

23. Svidritskiy, E. & Korostelev, A. A. Mechanism of Inhibition of Translation Termination by Blasticidin S. J. Mol. Biol. 430, 591–593 (2018).

24. Svidritskiy, E., Ling, C., Ermolenko, D. N. & Korostelev, A. A. Blasticidin S inhibits translation by trapping deformed tRNA on the ribosome. Proc. Natl. Acad. Sci. 110, 12283–12288 (2013).

25. Polikanov, Y. S. et al. Distinct tRNA Accommodation Intermediates Observed on the Ribosome with the Antibiotics Hygromycin A and A201A. Mol. Cell 58, 832–844 (2015).

26. Vázquez-Laslop, N. & Mankin, A. S. How Macrolide Antibiotics Work. Trends Biochem. Sci. 43, 668–684 (2018).

27. Beckert, B. et al. Structural and mechanistic basis for translation inhibition by macrolide and ketolide antibiotics. Nat. Commun. 12, 4466 (2021).

28. Anderson, M. G., Khoo, C. L.-Y. & Rickards, R. W. OXIDATION PROCESSES IN THE BIOSYNTHESIS OF THE TETRACENOMYCIN AND ELLORAMYCIN ANTIBIOTICS. J. Antibiot. (Tokyo) 42, 640–643 (1989).

29. Drautz, H., Reuschenbach, P., Zähner, H., Rohr, J. & Zeeck, A. Metabolic products of microorganisms. 225. Elloramycin, a new anthracycline-like antibiotic from Streptomyces olivaceus. Isolation, characterization, structure and biological properties. J. Antibiot. (Tokyo) 38, 1291–1301 (1985).

30. Egert, E., Noltemeyer, M., Siebers, J., Rohr, J. & Zeeck, A. THE STRUCTURE OF TETRACENOMYCIN C. J. Antibiot. (Tokyo) 45, 1190–1192 (1992).

31. Rohr, J. & Zeeck, A. Structure-activity relationships of elloramycin and tetracenomycin C. J. Antibiot. (Tokyo) 43, 1169–1178 (1990).

32. Weber, W., Zähner, H., Siebers, J., Schröder, K. & Zeeck, A. [Metabolic products of microorganisms. 175. Tetracenomycin C (author’s transl)]. Arch. Microbiol. 121, 111–116 (1979).

33. Guillerez, J., Lopez, P. J., Proux, F., Launay, H. & Dreyfus, M. A mutation in T7 RNA polymerase that facilitates promoter clearance. Proc. Natl. Acad. Sci. U. S. A. 102, 5958–5963 (2005).

34. Zhang, J., Kobert, K., Flouri, T. & Stamatakis, A. PEAR: a fast and accurate Illumina Paired-End reAd mergeR. Bioinformatics 30, 614–620 (2014).

35. Love, M. I., Huber, W. & Anders, S. Moderated estimation of fold change and dispersion for RNA-seq data with DESeq2. Genome Biol. 15, 550 (2014).

36. Hunter, J. D. Matplotlib: A 2D Graphics Environment. Comput. Sci. Eng. 9, 90–95 (2007).

37. Tareen, A. & Kinney, J. B. Logomaker: beautiful sequence logos in Python. Bioinformatics 36, 2272–2274 (2020).

38. Tikhonova, E. B. & Zgurskaya, H. I. AcrA, AcrB, and TolC of Escherichia coli Form a Stable Intermembrane Multidrug Efflux Complex *. J. Biol. Chem. 279, 32116–32124 (2004).

39. Weston, N., Sharma, P., Ricci, V. & Piddock, L. J. V. Regulation of the AcrAB-TolC efflux pump in Enterobacteriaceae. Res. Microbiol. 169, 425–431 (2018).

40. New tools for automated cryo-EM single-particle analysis in RELION-4.0 - PubMed. https://pubmed.ncbi.nlm.nih.gov/34783343/.

41. Zheng, S. Q. et al. MotionCor2: anisotropic correction of beam-induced motion for improved cryo-electron microscopy. Nat. Methods 14, 331–332 (2017).

42. Rohou, A. & Grigorieff, N. CTFFIND4: Fast and accurate defocus estimation from electron micrographs. J. Struct. Biol. 192, 216–221 (2015).

43. Pettersen, E. F. et al. UCSF Chimera--a visualization system for exploratory research and analysis. J. Comput. Chem. 25, 1605–1612 (2004).

44. Terwilliger, T. C., Sobolev, O. V., Afonine, P. V. & Adams, P. D. Automated map sharpening by maximization of detail and connectivity. Acta Crystallogr. Sect. Struct. Biol. 74, 545–559 (2018).

45. Emsley, P. & Cowtan, K. Coot: model-building tools for molecular graphics. Acta Crystallogr. D Biol. Crystallogr. 60, 2126–2132 (2004).

46. Croll, T. I. ISOLDE: a physically realistic environment for model building into low-resolution electron-density maps. Acta Crystallogr. Sect. Struct. Biol. 74, 519–530 (2018).

47. Adams, P. D. et al. PHENIX: a comprehensive Python-based system for macromolecular structure solution. Acta Crystallogr. D Biol. Crystallogr. 66, 213–221 (2010).

48. Pettersen, E. F. et al. UCSF ChimeraX: Structure visualization for researchers, educators, and developers. Protein Sci. Publ. Protein Soc. 30, 70–82 (2021).

