## Supplementary figures and images for "Tetracenomycin X sequesters peptidyl-tRNA during translation of QK motifs"

### Source Data for Fig. 2a

##### Source Data for Fig. 2a

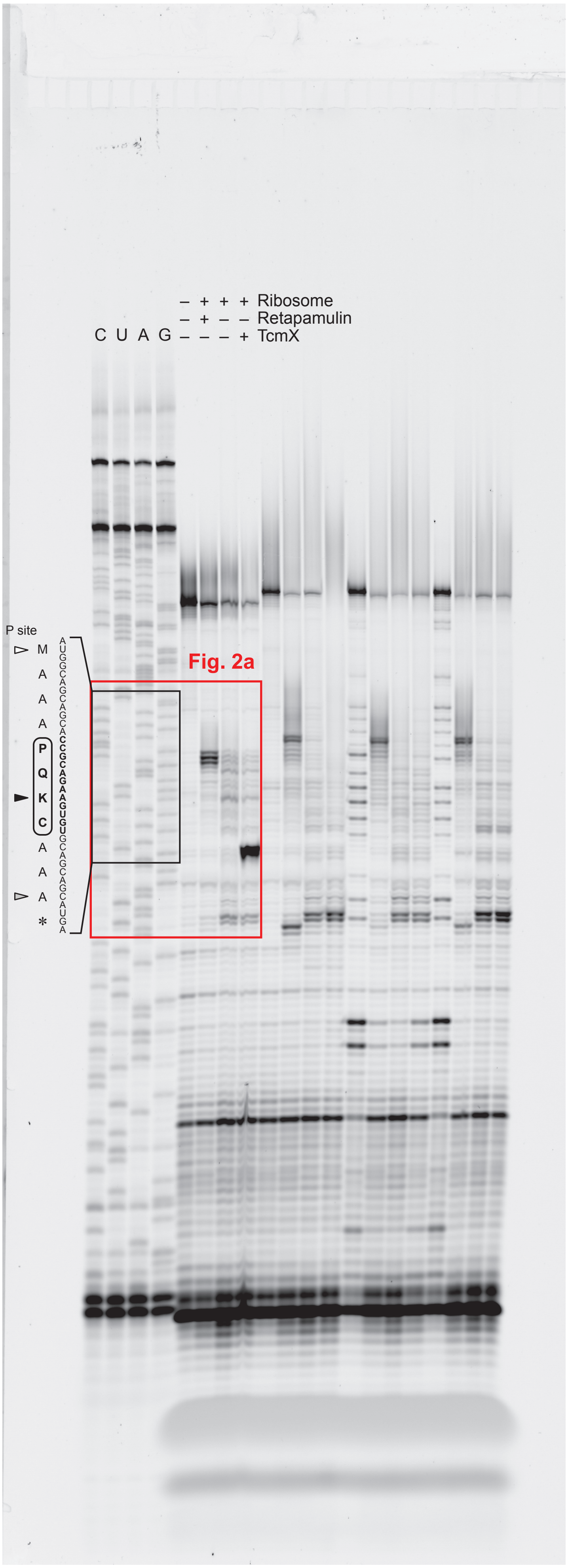

### Source Data for Fig. 2b

#### Source Data for Fig. 2b

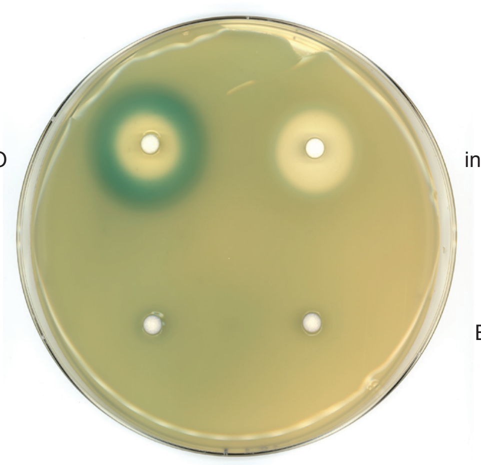


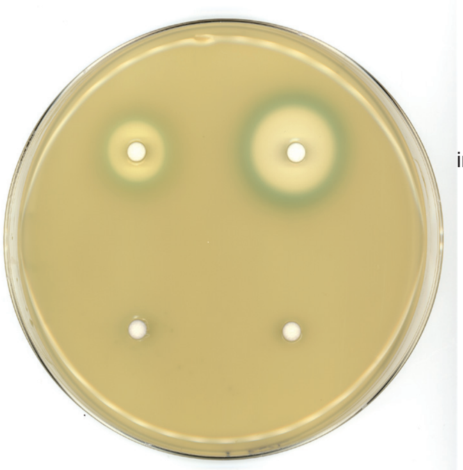
